# Evolutionary dynamics of piRNA clusters in *Drosophila*

**DOI:** 10.1101/2021.08.20.457083

**Authors:** Filip Wierzbicki, Robert Kofler, Sarah Signor

## Abstract

Small RNAs produced from transposable element (TE) rich sections of the genome, termed piRNA clusters, are a crucial component in the genomic defense against selfish DNA. In animals it is thought the invasion of a TE is stopped when a copy of the TE inserts into a piRNA cluster, triggering the production of cognate small RNAs that silence the TE. Despite this importance for TE control, little is known about the evolutionary dynamics of piRNA clusters, mostly because these repeat rich regions are difficult to assemble and compare. Here we establish a framework for studying the evolution of piRNA clusters quantitatively. Previously introduced quality metrics and a newly developed software for multiple alignments of repeat annotations (Manna) allow us to estimate the level of polymorphism segregating in piRNA clusters and the divergence among homologous piRNA clusters. By studying 20 conserved piRNA clusters in multiple assemblies of four *Drosophila* species we show that piRNA clusters are evolving rapidly. While 70-80% of the clusters are conserved within species, the clusters share almost no similarity between species as closely related as *D. melanogaster* and *D. simulans*. Furthermore, abundant insertions and deletions are segregating within the *Drosophila* species. We show that the evolution of clusters is mainly driven by large insertions of recently active TEs, and smaller deletions mostly in older TEs. The effect of these forces is so rapid that homologous clusters often do not contain insertions from the same TE families.x

## Introduction

Transposable elements (TEs) are short sequences of DNA that multiply within genomes [McClintock, 1956]. TEs are widespread across the tree of life, often making up a significant portion of the genome (2.7-25% in fruit flies, 45% in humans, and 85% in maize [Piegu et al., 2006, Schnable et al., 2009, Lee and Langley, 2012]). TEs also impose a severe mutational load on their hosts by producing insertions that disrupt functional sequences and mediate ectopic recombination [Lim, 1988, Levis et al., 1984, McGinnis et al., 1983]. However, some TE insertions have also been associated with increases in fitness, for example due to changes in gene regulation, where they can act as enhancers, repressors, or other regulators of complex gene expression patterns [Daborn et al., 2002, González et al., 2008, Mateo et al., 2014, Casacuberta and González, 2013]. The distribution of fitness effects of TEs is not known, but the majority of insertions are thought to be deleterious [Yang and Nuzhdin, 2003, Dimitri et al., 2003, Lee and Langley, 2012, Adrion et al., 2017].

For a long time TEs were thought to be solely counteracted at the population level (transposition/selection balance) [Charlesworth and Charlesworth, 1983, Barrón et al., 2014]. However the discovery of a small RNA-based defense system revealed that they are also actively combated by the host [Brennecke et al., 2007, Lee and Langley, 2010, Blumenstiel, 2011]. This host defense system relies upon PIWI interacting RNAs (piRNAs) that bind to PIWI-clade proteins and suppress TE activity transcriptionally and post-transcriptionally [Brennecke et al., 2007, Gunawardane et al., 2007, Sienski et al., 2012, Le Thomas et al., 2013]. For example in *D. melanogaster* post-transcriptional silencing of TEs is based on Aub and Ago3 which, guided by piRNAs, cleave TE transcripts in the cytoplasm [Kalmykova et al., 2005, Peters and Meister, 2007, Brennecke et al., 2007, Gunawardane et al., 2007]. In the nucleus piRNAs guide the Piwi protein to transcribed TEs which, aided by other proteins, transcriptionally silence TEs through the establishment of repressive chromatin marks [Sienski et al., 2012, Le Thomas et al., 2013, Darricarrere et al., 2013]. These piRNAs are produced from discrete regions of the genome termed piRNA clusters, which largely consist of TE fragments [Brennecke et al., 2008]. There is evidence that a single insertion of a TE into a piRNA cluster may be sufficient to initiate piRNA mediated silencing of the TE [Marin et al., 2000, Josse et al., 2007, Zanni et al., 2013]. Therefore, it is assumed that a newly invading TE proliferates in the host until a copy jumps into a piRNA cluster, which triggers the production of piRNAs that silence the TE [Bergman et al., 2006, Malone and Hannon, 2010, Goriaux et al., 2014, Ozata et al., 2019].

Despite the central importance of piRNA clusters for the control of TEs, we know very little about how piRNA clusters evolve within and between species. For example, transposition into clusters would be advantageous to hosts if cluster insertions are indeed required for functional silencing of TEs. Then, a general expansion of piRNA clusters would be expected with the invasion of novel TEs. Such invasions may be quite frequent. For example it is likely that four TE families invaded worldwide *D. melanogaster* populations within the last 100years [Schwarz et al., 2021]. Larger or more abundant piRNA clusters in turn will expand the functional target for transposition and may thus be favored. In support of this hypothesis it was suggested that piRNA clusters have largely been gained over the course of evolution [Chirn et al., 2015]. However, these claims are difficult to evaluate as studying the evolution of piRNA clusters is challenging for several reasons. First, piRNA clusters are highly repetitive and very difficult to assemble, thus high quality ungapped assemblies of these repetitive regions are required [see for example Wierzbicki et al., 2021] Second, it is challenging to unambiguously identify homologous clusters within and between species. Third, investigating the evolution of the composition of clusters requires reliable alignments of the highly repetitive piRNA clusters. Due to these challenges and the importance of these clusters for TE control, the evolutionary turnover of piRNA clusters is considered to be a central open question in TE biology [Czech et al., 2018].

Here, we investigate the evolution of piRNA clusters within and between four *Drosophila* species. By combining long-read based assemblies with a recently developed approach for identifying homologous piRNA clusters (CUSCO, [Wierzbicki et al., 2021]) and a newly developed software for generating multiple alignments of repetitive regions (Manna) we are able to shed light on the evolution of piRNA clusters. While piRNA clusters are 70-80% conserved within species, they share almost no similarity between species as closely related as *D. melanogaster* and *D. simulans*. Many polymorphic insertions and deletions within clusters are maintained in *Drosophila* populations. The evolutionary forces dictating the observed patterns appear to be large insertions of recently active TEs, and smaller deletions of older TE insertion. Due to this rapid turnover, homologous piRNA clusters frequently do not contain insertions from the same TE families. Using our approach of combining CUSCO and Manna, we established a framework to study piRNA cluster evolution quantitatively within and between species.

## Results

### Identification of homologous piRNA clusters

To shed light on the evolution of piRNA clusters, we compared the composition of clusters among related *Drosophila* species. *D. sechellia, D. mauritiana*, and *D. simulans* are closely related, having an estimated divergence time of 0.7 million years, while *D. melanogaster* diverged from this group 1.4 million years ago (fig. 1A, [Obbard et al., 2012]). We relied on long-read assemblies as they allow for end to end reconstruction of piRNA clusters and their TE content and thus promise to provide a complete picture of cluster evolution [Wierzbicki et al., 2021]. Since we are interested in the evolution of clusters both within and between species, we obtained long-read assemblies of several strains for *D. melanogaster* and *D. simulans*. In total we analyzed nine long-read based assemblies, four of *D. simulans*, three of *D. melanogaster* , and one each of *D. sechellia* and *D. mauritiana*. Seven assemblies were publicly available and two assemblies of *D. simulans* strains were generated in this work with Oxford Nanopore long reads (*SZ45, SZ232*) [Chakraborty et al., 2021, Nouhaud, 2018, Signor et al., 2017a].

**Figure 1:**
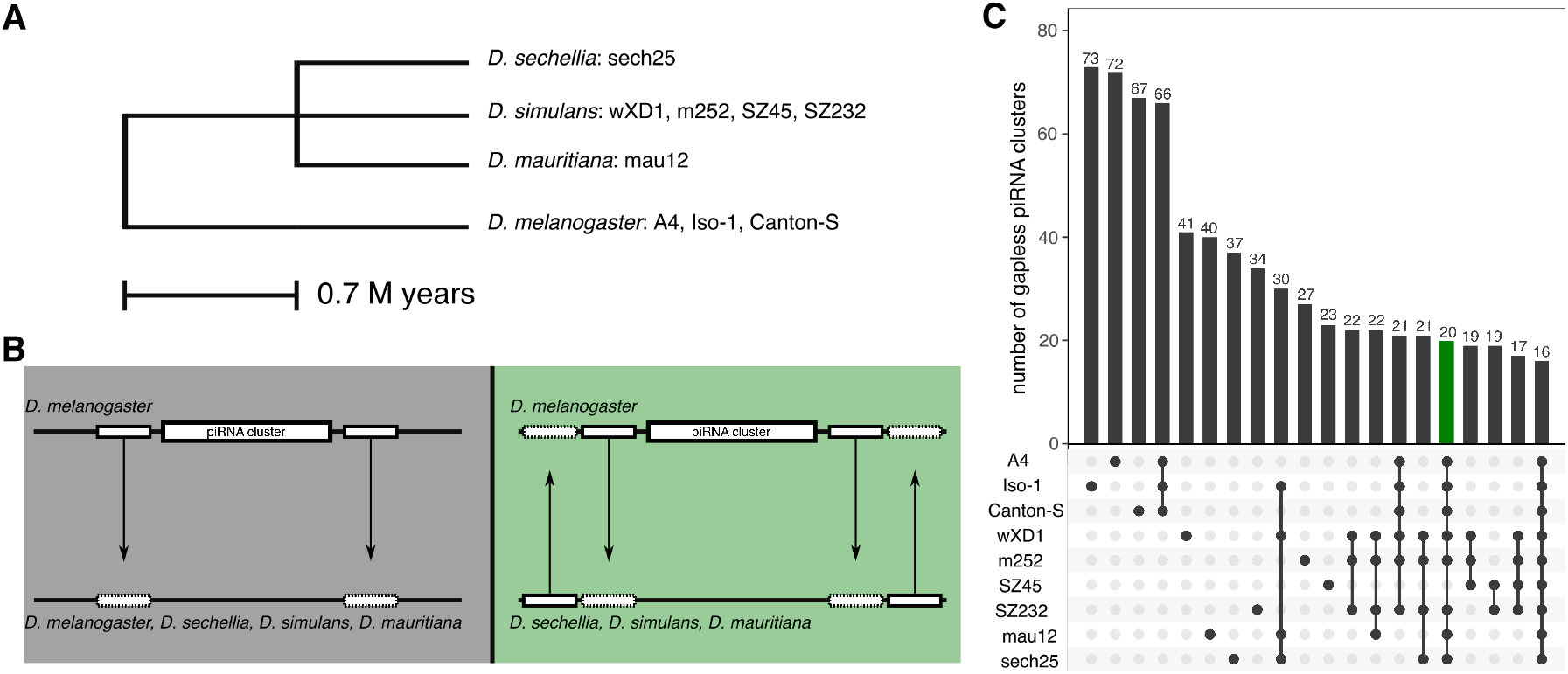
Overview of the species and piRNA clusters used in this work. A) Phylogenetic tree showing the evolutionary distance between the four species investigated in this work (based on [Obbard et al., 2012]). The analyzed strains are shown after the species name. B) Our approach for finding homologous piRNA clusters in the different species and strains. Unique sequences flanking piRNA clusters were aligned to the target strain. An homologous cluster was identified when both flanking sequences aligned to the same contig (grey). We confirmed homology of clusters by designing flanking sequences in the target strain and aligning them back to *D. melanogaster* reference genome (green, “reciprocal flanks”). C) Number of gapless piRNA clusters found in different species/strains. Colors of bar (grey or green) correspond to the approach used for identifying homologous clusters (see B)

The identification of homologous piRNA clusters among the different strains and species was based on unique sequences flanking 85 out of the 142 piRNA clusters in *D. melanogaster* (flanking sequences could not be designed for telomeric clusters extending to the ends of chromosomes or clusters on the fragmented U-chromosome) [Wierzbicki et al., 2021]. These flanking sequences were mapped to each assembly, and homologous piRNA clusters were identified as the regions between the aligned flanking sequences (fig. 1B; grey). piRNA clusters with assembly gaps or flanking sequences aligning to different contigs were not considered. To validate the homology of the piRNA clusters, we designed additional pairs of flanking sequences in the target species, aligned them back to *D. melanogaster* and ascertained that these mapped sequences flank the piRNA clusters of *D. melanogaster* (fig. 1B,C; green; supplementary tables S1-S3). The number of assembled piRNA clusters varied considerably between the strains and species, ranging from 73 clusters in *D. melanogaster Iso-1* to 23 clusters in *D. simulans SZ45* (fig. 1C). To study the evolution of piRNA clusters between species, we focused on 20 piRNA clusters shared between *D. mauritiana, D. sechellia* and the three best assemblies of *D. melanogaster* and *D. simulans* (fig. 1C; red). Most notably our analysis included clusters *42AB* (cluster 1), *20A* (cluster 2) and *38C* (cluster 5) but not *flamenco*. Except for cluster *20A*, which is an uni-strand cluster that is expressed in the germline and the soma, all analyzed clusters are dual-strand clusters that are solely expressed in the germline [Mohn et al., 2014, Brennecke et al., 2007]. By investigating the heterogeneity of the base coverage and the softclip coverage - two recently proposed metrics for identifying assembly errors in piRNA clusters [Wierzbicki et al., 2021] - we ascertained that the assemblies of the 20 clusters are of high quality (see Materials and Methods; supplementary figs. S1-S5). Based on publicly available small RNA data from ovaries of a *D. melanogaster* and *D. simulans* strain collected in Chantemesle (France; [Asif-Laidin et al., 2017]), we found that 15 out of the 20 investigated clusters are expressed in both species (*>* 10 reads per million; supplementary figs. S6, S7, S8).

### Comparing the composition of homologous clusters

piRNA clusters are often referred to as ‘TE graveyards’ since they are thought to carry the remains of past TE invasions. This highly repetitive nature makes it difficult to compare the composition of homologous clusters, e.g. using multiple sequence alignments. We approached this problem inspired by the alignments of amino-acid sequences, which are performed at a higher level than the underlying nucleotide sequences. Here, we propose that multiple alignments may be performed with the TE annotations (e.g. generated by RepeatMasker) of piRNA clusters instead of the nucleotide sequences. For this reason, we developed Manna (<u>m</u>ultiple <u>ann</u>otation <u>a</u>lignment), a novel tool performing multiple alignments of annotations. Although primarily designed for annotations of repeats, it may work with the annotations of any feature. Manna performs a progressive alignment similar to that described by [Feng and Doolittle, 1987]. Using a simple scoring scheme (supplementary fig. S9) and an adapted Needleman-Wunsch algorithm [Needleman and Wunsch, 1970] a guide tree is computed. Based on this tree the most similar annotations are aligned first, followed by increasingly more distant annotations. For the scoring matrix the score of each newly aligned annotation is computed as the average score of the previously aligned annotations [Feng and Doolittle, 1987].

This novel tool enables us to compare the composition of homologous clusters using the following approach: First, we align pairs of sequences flanking piRNA clusters to the assemblies, thereby identifying the positions of homologous clusters in each assembly (fig. 2A). Second, we extract the sequences delimited by these pairs of flanking sequences (fig. 2B). Third, we annotate repeats in the extracted sequences (fig. 2C) and solely retain the repeat annotation (fig. 2D). Finally, we align the repeat annotation with Manna (fig. 2E). Using simulated sequences with varying repeat contents, we carefully validated this approach for comparing the composition of homologous piRNA clusters (supplementary results S1).

**Figure 2:**
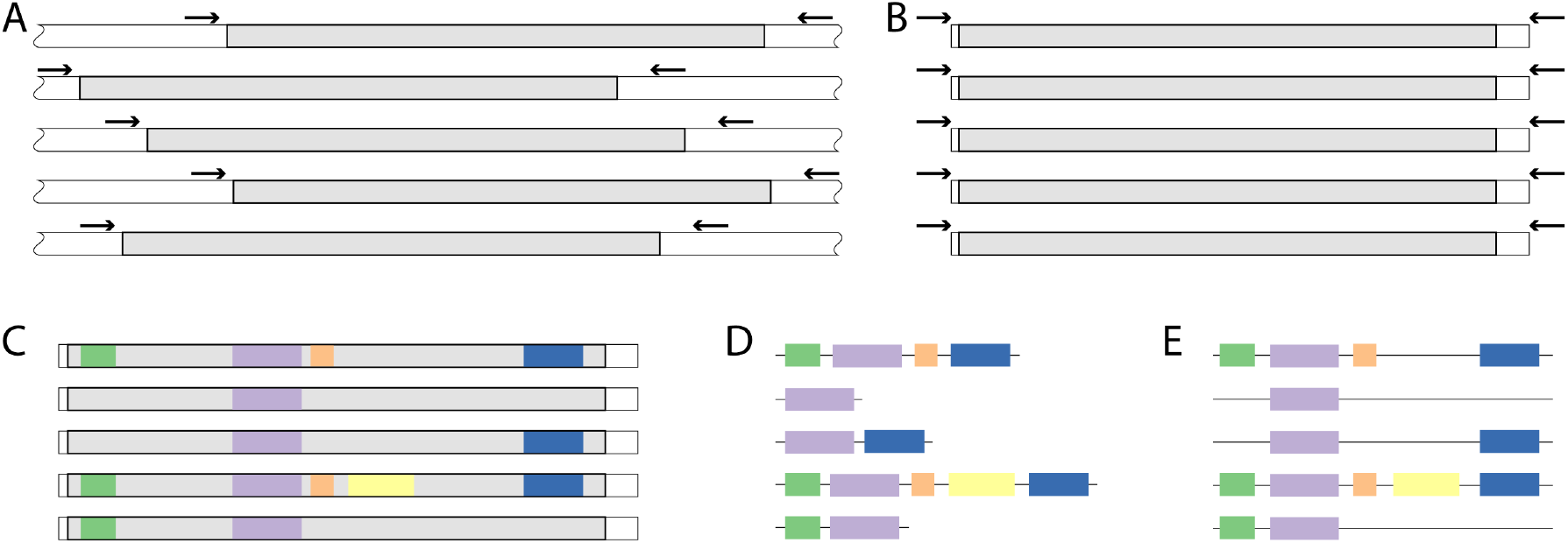
Overview of our approach for comparing the composition of piRNA clusters. A) To identify homologous piRNA clusters (grey areas) in the strains, we mapped sequences flanking the piRNA clusters (black arrows) to the assemblies. B) Regions delimited by the flanking sequences were extracted (i.e. the piRNA clusters plus the short sequences between the clusters and the flanking sequences). C) Repeats were annotated in the extracted sequences. D) Solely the repeat annotations were retained for further analysis. E) The repeat annotations were aligned with Manna allowing us to compare the repeat content of piRNA clusters.

Alignments with Manna allow us to quantify i) the number of polymorphic and fixed TE insertions and ii) the similarity *s* and the distance (*d* = 1 *– s*) among homologous clusters. The similarity (*s*) between clusters is computed as *s* = 2 *\* a/*(2 *\* a* + *u*) where *a* and *u* are the total length of aligned and unaligned TE sequences, respectively (for examples see supplementary fig. S10). This similarity can be intuitively interpreted as the fraction of TE sequences that can be aligned between two (homologous) clusters.

Alignments with Manna do not incorporate unannotated sequence in between TEs (fig. 2C). Therefore, we additionally investigated the similarity among homologous clusters using a complementary approach: we identified similar sequences between clusters with BLAST (minimum identity 70% [Altschul et al., 1990]) and visualized these similarities and the repeat content of clusters with Easyfig (supplementary figs. S11-S15).

### Rapid evolution of piRNA clusters

To quantify the rate at which piRNA clusters evolve, we estimated the evolutionary turnover of the TE content of the 20 piRNA clusters using the similarity (*s*) as computed with Manna (see above). Based on the distance between the clusters (*d* = 1 *– s*), we additionally generated phylogenetic trees reflecting these distances (fig. 3A).

**Figure 3:**
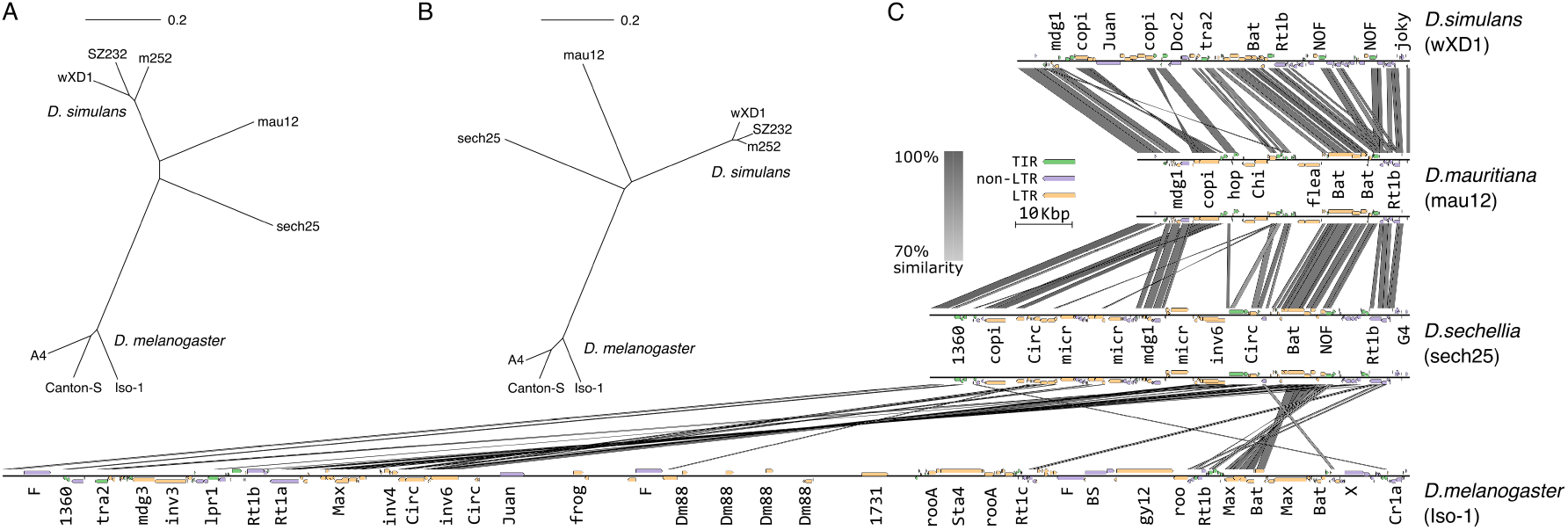
piRNA clusters are rapidly evolving in *Drosophila* species. A) Phylogenetic tree summarizing the distance of the 20 piRNA clusters among the different strains and species weighted by the average cluster lengths. The distance is estimated by Manna as the fraction of unaligned TE sequences (scale bar shows a distance of 20%). Note that solely about 8.1% of the TE sequences can be aligned between the clusters of *D. melanogaster* (green) and *D. simulans* (blue). B) Phylogenetic tree for the piRNA cluster 42AB (cluster 1) based on alignments with Manna. C) The evolution of piRNA cluster *42AB* in four *Drosophila* species visualized with Easyfig. Homology among the sequences (grey bars) was determined with BLAST. The grey scale indicates the degree of the sequence similarity. Homology blocks smaller than 400bp are not shown. Insertions of TEs are shown as small rectangular arrows where the color indicates the order (LTR, non-LTR and TIR). Family names are abbreviated.

Strikingly, an average of solely 8.1% of the TE sequences can be aligned between the piRNA clusters of *D. melanogaster* and *D. simulans* (fig. 3A; supplementary table S4). Among the 20 clusters the similarity ranged from 0.0% for clusters 19 and 110 to 93.5% for cluster 114 (length weighted median: 3.7%; supplementary table S4). Within the more closely related species of the *simulans* complex 41.4% of the TE sequences can be aligned between *D. simulans* and *D. mauritiana* (range: 0.0 - 100%; length weighted median: 32.7%) and 32.7% between *D. sechellia* and *D. simulans* (range: 0.0 - 88.8%; length weighted median: 24.8%; supplementary table S4). Our data thus suggest that the clusters of *D. simulans* are more closely related to *D. mauritiana* than to *D. sechellia*. Given this rapid turnover within piRNA clusters, we also hypothesized that there should be abundant polymorphisms within species. In agreement with this, we found that the average similarity of clusters within species is 73.12% for *D. melanogaster* (range: 33.3-100%; length weighted median: 74.2%) and 74.7% for *D. simulans* (range: 0.0-100%; length weighted median: 75%; supplementary table S4). That is to say that on average 26% of the TE sequences in piRNA clusters cannot be aligned between two assemblies of the same species. The TE content of clusters is thus highly polymorphic within species.

However, the strains analyzed in *D. simulans* and *D. melanogaster* were collected at very diverse time points and geographic locations. We therefore speculated that the similarity among strains sampled from the same population may be higher. A comparison of 16 clusters shared between the Californian *D. simulans* strains *SZ232* and *SZ45*, which were collected at the same location and date, an African strain (*m252*) and an old Californian strain (*w*^*xD*1^, likely collected approximately 50 years prior) did not confirm this hypothesis (similarity between *SZ232* vs. *SZ45* : 72.5%; average similarity among all other *D. simulans* strains: 75.8%; supplementary table S5). The clusters of strains sampled from the same population are thus not necessarily more similar than the clusters of strains sampled from different regions and time points (although the results vary among the clusters).

Next, we aimed to investigate the evolution of cluster *42AB* (cluster 1) in more detail. In *D. melanogaster 42AB* is one of the largest clusters that may account for 20-30% of all piRNAs [Brennecke et al., 2007]. It is thus frequently highlighted as a canonical piRNA cluster [e.g. Czech et al., 2008, Mohn et al., 2014, Olovnikov et al., 2013, Andersen et al., 2017]. A phylogenetic tree based on an alignment of annotated TEs shows that cluster *42AB* is rapidly evolving among the investigated *Drosophila* species (fig. 3B; for a tree for all other clusters see supplementary fig. S16). The similarity of *42AB* between *D. simulans* and *D. melanogaster* , based on an alignment of TE annotations using Manna, is solely 4%. Within the *simulans* clade the similarity of *42AB* between *D. simulans* and *D. mauritiana* is 29.6%, and between *D. simulans* and *D. sechellia* it is 26.4% (supplementary table S4). Within species, cluster *42AB* is more variable in *D. melanogaster* (similarity: 77.5%) than in *D. simulans* (similarity: 90.3% ; supplementary table S4). As alignments with Manna only capture similarities of annotated TEs we also visualized the evolution of cluster *42AB* using BLAST and Easyfig (fig. 3C). This approach confirms our findings. Cluster *42AB* has few sequence similarities between *D. melanogaster* and *D. simulans* and a higher level of sequence similarity among the species of the *simulans* complex (fig. 3C). We conclude that cluster *42AB* is rapidly evolving in the investigated species (fig. 3C). For a visualization of the sequence similarity of all 20 clusters in the four species see supplementary figs. S11-S15.

Thus far we have shown that the sequence of piRNAs clusters is evolving very quickly between and within species. However, it is possible that this rapid evolution is due to rearrangements within piRNA clusters [Gebert et al., 2021], while the TE content of clusters actually remains stable. We addressed this question by quantifying the number of insertions from each TE family in each cluster, and determining if at least one insertion of a given family is present in a given cluster in *D. simulans, D. melanogaster* or both species (an insertion in any of the three strains of each species was considered as a presence). For example we considered *blood* to be present in cluster *42AB* in both species when a single *blood* insertion was found in *42AB* of *A4* (*D. melanogaster*) and *m252* (*D. simulans*) but not in any other strain of the two species. The rapid evolution of piRNA clusters does not appear to be due to rearrangements, as the presence of TE families was also not conserved across species (fig. 4). Out of 321 TE families in piRNA clusters, only 76 were present in both species (families present in more than one cluster were counted multiple times). 164 were private to *D. melanogaster* and 81 to *D. simulans* (fig. 4). A similar observation can be made when we compare the TE composition of piRNA clusters among *D. simulans, D. mauritiana*, and *D. sechellia* (supplementary fig. S17).

**Figure 4:**
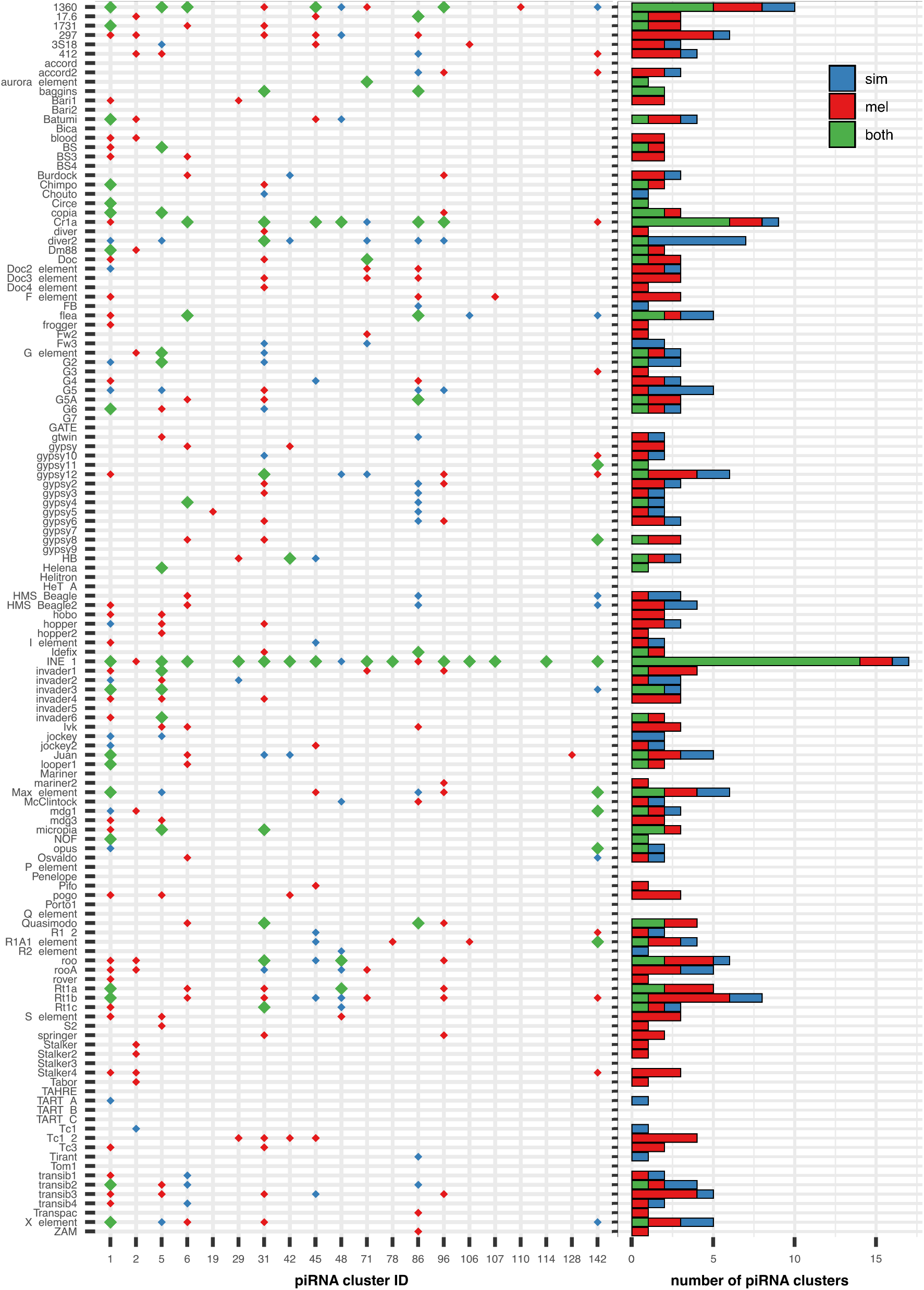
Overview of the TE content of piRNA clusters in *D. simulans* and *D. melanogaster* . For each piRNA cluster (x-axis) we indicate whether a given TE family (y-axis) has at least one insertion in *D. melanogaster* (red), *D. simulans* (blue) or in both species (green). We considered insertions in any of the three assemblies of *D. melanogaster* and *D. simulans*. The right panel summarizes the abundance of the families in piRNA clusters. Note that the TE content of the clusters varies dramatically between the species.

We thus conclude that piRNA clusters are rapidly evolving in *Drosophila* species, such that the average, only about 8% of TEs sequences can be aligned between the closely related *D. melanogaster* and *D. simulans*. Furthermore, homologous clusters frequently contain different TE families.

### piRNA clusters in *D. melanogaster* and *D. simulans* genotypes

Next, we investigated variation in the piRNA clusters of *D. melanogaster* and *D. simulans* in more detail, incorporating several genotypes from each species. An alignment of the 20 clusters with Manna in the three strains of *D. melanogaster* and *D. simulans* shows that clusters in *D. melanogaster* contain more TEs than in *D. simulans* (*Dmel* = 1, 002, *Dsim* = 547). The majority of these insertions are fixed (*Dmel* = 647, *Dsim* = 362; fig. 5A), but a substantial number of TE insertions is segregating in one (*Dmel* = 229, *Dsim* = 118) or two genotypes (*Dmel* = 126, *Dsim* = 67). Despite these differences in the TE abundance among the two species, the site frequency spectrum of the cluster insertions is very similar between *D. melanogaster* and *D. simulans* (Chi-squared test *p* = 0.20; fig. 5A). The large number of polymorphic cluster insertions is not contingent upon a single outlier-genotype since all genotypes from both species carried abundant polymorphic cluster insertions (*D. melanogaster* : *CS* = 191, *A*4 = 153, *Iso*1 = 137; *D. simulans SZ*232 = 106, *w*^*xD*1^ = 97, *m*252 = 49, supplementary fig. S18A). The polymorphic cluster insertions were distributed over 17 clusters in *D. melanogaster* and 12 clusters in *D. simulans* (supplementary fig. S18A). In agreement with the higher TE content of *D. melanogaster* clusters, piRNA clusters in *D. melanogaster* were substantially longer than in *D. simulans* (Wilcoxon rank sum test *W* = 2192, *p* = 0.040; supplementary fig. S18B). The total size of the piRNA clusters in *D. melanogaster* was about double that of the clusters in *D. simulans* (average over all three strains *dmel* = 817, 770, *dsim* = 452, 591). In both species segregating cluster insertions were on the average longer than fixed ones (*D. melanogaster* : *seg* = 1115, *fix* = 591, Wilcoxon rank sum test *W* = 122302, *p* = 0.089; *D. simulans*: *seg* = 798, *fix* = 470, Wilcoxon rank sum test *W* = 38248, *p* = 0.0065).

**Figure 5:**
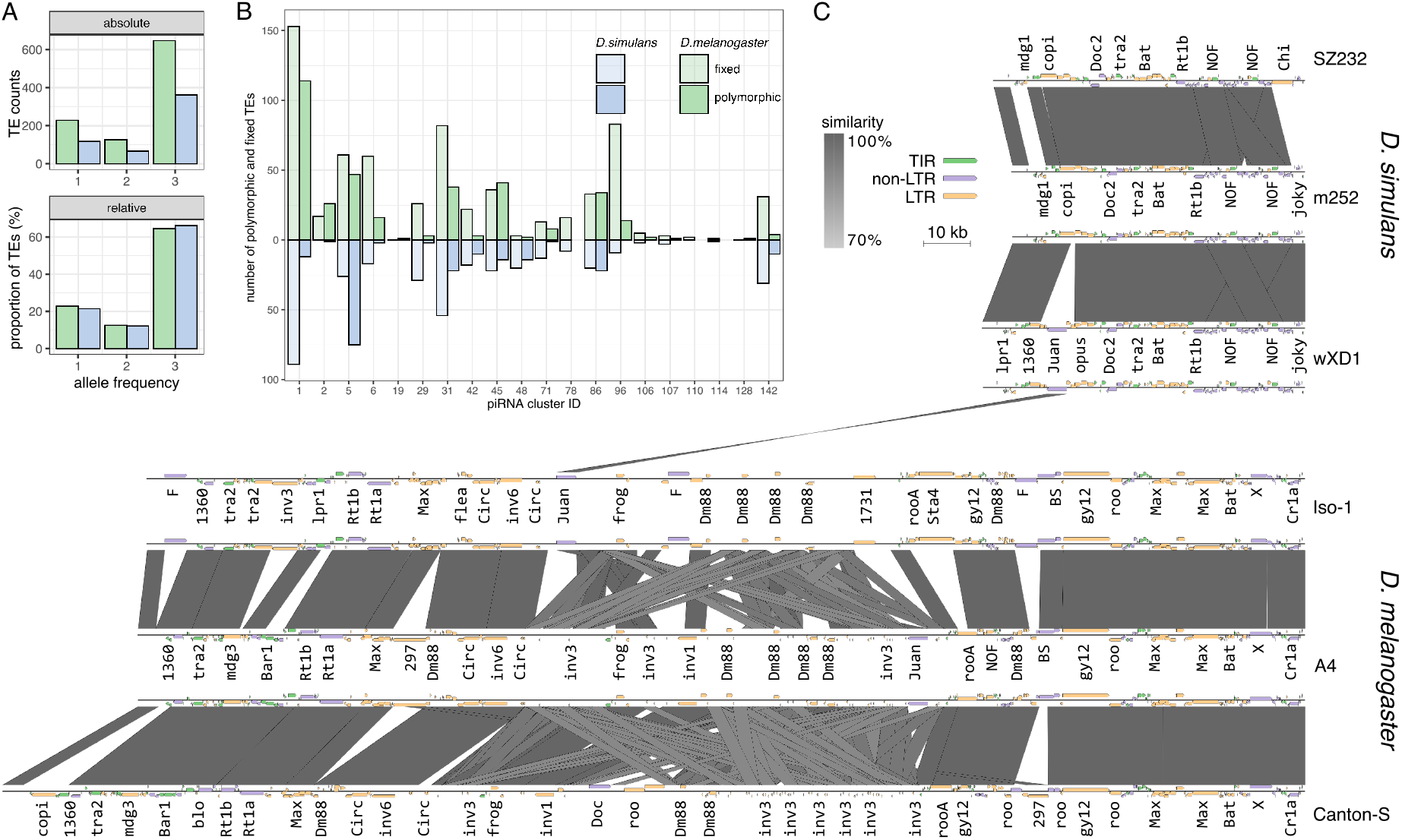
Rapid evolution of piRNA clusters within *D. melanogaster* and *D. simulans*. A) Population frequencies of TE insertions in all 20 piRNA clusters of *D. melanogaster* (green) and *D. simulans* (blue). The absolute (top) and relative (bottom) TE abundance are shown. Insertions occurring in three individuals are fixed. B) Numbers of fixed (transparent) and polymorphic (opaque) sites for each cluster in *D. melanogaster* (green) and *D. simulans* (blue). C) Composition of cluster 42AB in 3 strains of *D. melanogaster* and *D. simulans*. Grey bars indicate regions of similarity among two assemblies of 42AB (minimum length 3 kb). TE families are colored by order (LTR, non-LTR and TIR).

In addition, the amount of polymorphism segregating in strains sampled from the same population (*SZ232, SZ45*) is similar to the amount of polymorphism sampled in strains from different locations (*m252*, Africa) and time points (*w*^*xD*1^, California; percent polymorphic insertions with a minimum size of 100bp: *SZ232* vs *SZ45* = 23.8%, mean of all other pairwise comparisons = 20.7%; supplementary figs. S19, S20). While overall polymorphism was similar amongst strains, the amount of fixed and segregating TE insertions varies across the clusters. Some clusters in *D. melanogaster* mostly have fixed TEs such as cluster 96 (*fix* = 83, *seg* = 14) and cluster 142 (*fix* = 31, *seg* = 4), but other clusters, like cluster 1 (*fix* = 153, *seg* = 114) and cluster 45 (*fix* = 36, *seg* = 41), have large proportions of segregating TEs (fig. 5B). Similarly in *D. simulans* some clusters such as cluster 1 (*fix* = 89, *seg* = 12) and cluster 29 (*fix* = 29, *seg* = 2) have largely fixed TEs whereas cluster 5 (*fix* = 26, *seg* = 75) and cluster 86 (*fix* = 20, *seg* = 22) contain many segregating TE insertions. This indicates that clusters may evolve at different rates, with some clusters evolving faster than others. Additionally, the evolutionary turnover of the clusters may differ among species, for example cluster *42AB* (cluster 1) evolves faster in *D. melanogaster* whereas cluster *5* evolves faster in *D. simulans* (fig. 5B).

Our analysis is based on the consensus sequences of *D. melanogaster* TEs. We asked if this could lead to a bias where TE insertions in *D. simulans* clusters are less readily identified than in *D. melanogaster* . Such a bias should lead to a lower density of TEs in piRNA clusters of *D. simulans* as compared to *D. melanogaster* . We found that the density of TE insertions in piRNA clusters is very similar in the two species (TE insertions per kb *dmel* = 0.994, *dsim* = 0.985) suggesting that we identified most TE insertions in *D. simulans*. However, cluster insertions in *D. simulans* were, on the average, slightly shorter than in *D. melanogaster* (average length *dmel* = 777, *dsim* = 581; Wilcoxon rank sum test *W* = 300760, *p* = 0.0015). This is in agreement with previous works suggesting that TEs in *D. simulans* are shorter than in *D. melanogaster* [Lerat et al., 2011, Vieira et al., 2012], but it could also be a technical artefact where parts of TEs are not annotated in D. simulans due to the divergence of the TE from the consensus sequences.

Finally, we investigated the composition of cluster *42AB* in more detail (fig. 5D). Cluster *42AB* is, consistently among the strains, shorter in *D. simulans* than in *D. melanogaster* (fig. 5D; supplementary fig. S18B). The density of TEs in cluster 42AB is higher in *D. simulans* (TEs per kb *dmel* = 0.79, *dsim* = 1.41) possibly due to the shorter TE insertions (average length of TEs in 42AB *dmel* = 920*bp, dsim* = 658*bp*). While there is considerable sequence conservation in both species the *D. melanogaster 42AB* cluster bears no resemblance to *42AB* in *D. simulans*, other than containing a *Juan* element which is likely not a homologous insertion (fig. 5B). The number of segregating insertions is larger in *D. melanogaster* than in *D. simulans* suggesting that *42AB* is evolving faster in *D. melanogaster* (fig. 5B,D). For a visualization of the sequence similarity of all clusters in the different assemblies of *D. melanogaster* and *D. simulans* see supplementary figs. S11-S15.

We conclude that piRNA clusters are highly polymorphic in both species, that clusters have a similar TE density in both species and that most clusters are shorter in *D. simulans* than in *D. melanogaster* . Furthermore, clusters may evolve at different rates among and within species.

### Evolutionary forces shaping the composition of piRNA clusters

Many diverse evolutionary forces may act on the TE content of piRNA clusters, such as mutations, insertion bias, negative or positive selection and drift [Kofler, 2019, Kelleher et al., 2018, Lu and Clark, 2010, Brennecke et al., 2007, Zhang et al., 2020]. While we cannot distinguish among these forces we can shed light on their joint effect by investigating the abundance of insertions and deletions segregating in piRNA clusters. We determined the number of insertions and deletions segregating in piRNA clusters of the *D. simulans* strains by polarizing segregating indels using *D. mauritiana* as outgroup. We used TE insertions with a minimum length of 100 bp and considered indels resulting from presence/absence polymorphisms in the alignment and indels resulting from length differences between aligned TEs sequences. We found that 33 deletions and 99 insertions are segregating in piRNA clusters of *D. simulans* (fig. 6A) These indels were distributed over 12 of the investigated 20 piRNA clusters (supplementary fig S21). Insertions were, on the average, longer than deletions (average length 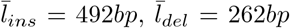; Wilcox rank sum test *W* = 920.5, *p* = 0.0002). Most indels were found in three of the 20 clusters: cluster 5 (43 indels), cluster 31 (20 indels), and cluster 45 (16 indels; supplementary fig. S21). Because *de novo* TE insertions will likely be large we separately analyzed long indels (*≥* 1000). We found that 12 long insertions and a single long deletion. The most abundant long insertions were due to the TE families *roo* and *Max-element* (two for each family). Both families are likely active in *D. simulans* [Kofler et al., 2015, Signor, 2020]. Finally, we asked if insertions are occurring with younger TE families than deletions. While we do not have direct estimates for the age of TE families in *D. simulans* we may use the average population frequency of all insertions of a family as proxy for age. Insertions of recently active families will mostly have a low frequency whereas old families will mostly have fixed insertions. Using the frequency estimates of Kofler et al. [2015] we found that families with insertions in piRNA clusters have a significantly lower average population frequency than families wit deletions (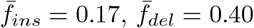; Wilcox rank sum test *W* = 2211, *p* = 2.7*e –* 05 fig. 6B).

**Figure 6:**
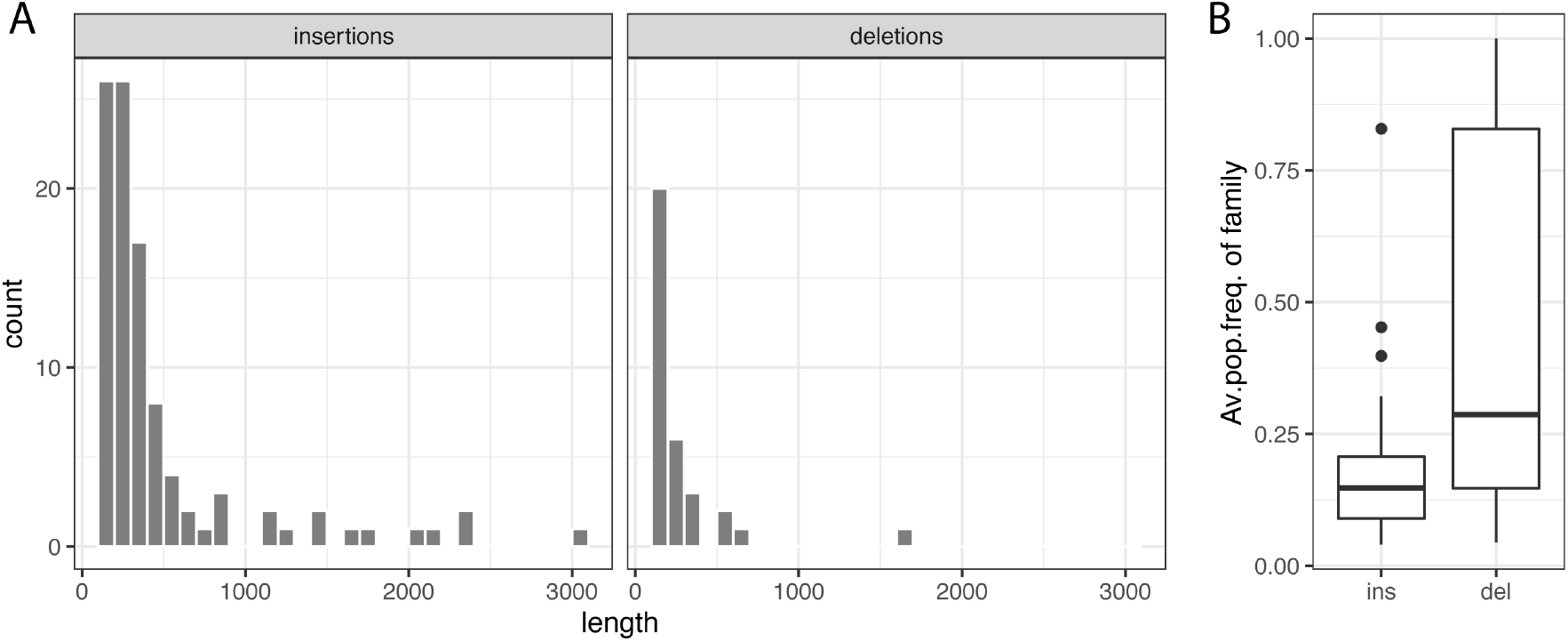
Overview of insertions and deletions in piRNA clusters of *D. simulans*. The clusters of *D. mauritiana* were used to polarize the indels. A) Histograms showing the abundance and length of insertions and deletions. B) Age of the families of insertions (ins) and deletions (del) in piRNA clusters, where the average population frequency (av.pop.freq.) of the family is used as a proxy for the age.

In summary, the evolutionary dynamics of piRNA clusters are governed by many insertions and few deletions, where insertions are on the average larger than deletions. Furthermore, insertions usually involve recently active families whereas deletions mostly happen in older families.

## Discussion

Here we established a framework for studying the evolution of piRNA clusters quantitatively, used that framework to analyze the composition of 20 piRNA clusters in four *Drosophila* species, and showed that piRNA clusters are evolving rapidly. This raises the question of whether the 20 piRNA clusters included in the analysis are a representative set of the 141 piRNA clusters of *D. melanogaster* . piRNA clusters were excluded from our analysis for three reasons i) clusters were at the end of a chromosome or on the unassembled U-chromosome which did not allow us to identify suitable flanking sequences ii) a cluster could not be assembled in all species without gaps, possibly due to complex repeat content iii) we could not identify conserved flanking sequences in all species such that the homology of a cluster could be established. While the first point likely does not introduce a bias the last two points could potentially result in a bias towards shorter or less complicated clusters. The analyzed clusters may thus be a rather conservative set, and it is possible that the excluded piRNA clusters have different evolutionary dynamics. To reduce possible biases in future works, it will be important to extend the analysis performed in the present work to a larger number piRNA clusters. It is possible that investigating alternate flanking sequences could lead to an increase in the number of clusters, and rapid advances in sequencing technology will increase the number of contiguously assembled clusters. However, a comparison between species will always be less than entirely comprehensive, as clusters may not be shared between species of interest or the flanking sequences may have degraded beyond recognition. In agreement with this, previous research has noted that many piRNA clusters are species specific [Gebert et al., 2021, Chirn et al., 2015].

This and other works established synteny of piRNA clusters based on sequences flanking the cluster up and downstream [Gebert et al., 2021, Chirn et al., 2015]. It is unclear if this is the best approach for finding homologous clusters. In principle, it is possible to use the sequence (or annotation) of piRNA clusters directly to search for the homologous clusters in an assembly of interest (e.g. with BLAST). However, given how rapidly piRNA clusters evolve, where solely 8% of TE sequences can be aligned between *D. melanogaster* and *D. simulans*, it is doubtful whether this approach will be able to correctly establish homology of the piRNA clusters. We quantified the similarity of clusters and the amount of polymorphism in clusters with our novel multiple alignment tool Manna. As a major innovation this tool performs a multiple alignment with repeat annotations rather than the raw sequences. While this approach provides invaluable insight into the evolution of piRNA clusters, it does ignore some information such as divergence of the TEs. Alignments of clusters at the nucleotide level may be more sensitive. But this approach has its own problems. Alignments of highly repetitive regions are challenging and may contain errors. Furthermore, the resulting alignment may be difficult to interpret. For example, it is unclear how to estimate the population frequency of a TE insertion where different parts of the TE align with several TE insertions in a homologous cluster. Manna avoids this fragmentation of TEs by aligning complete chunks of annotated TEs.

We found that *D. simulans* has fewer TE insertions in piRNA clusters than *D. melanogaster* . That this is a real pattern is supported by the similar density of TEs in the two species within the piRNA clusters (indicating no obvious presence of unannotated TEs in *D. simulans*). However, the TE libraries used here are curated to represent few overlapping TE families. It is still possible that in *D. simulans* some TEs are only partially annotated or missed entirely. If this were the case, then piRNA clusters in *D. simulans* would be denser than in *D. melanogaster* .

It is an important question which evolutionary forces drive the evolution of piRNA clusters. In principle, the following forces could act on piRNA clusters. First, different types of mutations, such as insertions due to recent TE activity, the deletion bias observed in *Drosophila* or major rearrangements, for example due to ectopic recombination mediated by TE insertions, may contribute to the rapid turnover of piRNA clusters [Petrov et al., 1996, Langley et al., 1988]. Many TE families are active in *Drosophila* species so recent insertions may be an important driver of cluster evolution [Kofler et al., 2015]. Also genomic rearrangements have been implicated in the evolution of clusters [Assis and Kondrashov, 2009, Gebert et al., 2021]. Second, selection (positive or negative) may contribute to the rapid evolution of piRNA clusters. Theory suggests that an invading TE is silenced by multiple segregating TE insertions distributed over many piRNA clusters [Kofler, 2019, Kelleher et al., 2018]. This hypothesis has been confirmed experimentally by recent works investigating the distribution of cluster insertions in natural and experimental populations that were recently invaded by a TE [Zhang et al., 2020, Kofler et al., 2018]. Theory further suggests that these segregating cluster insertions could be positively selected as haplotypes with a cluster insertion will accumulate few TEs overall and will thus be less deleterious than haplotypes without a cluster insertion [Kofler, 2019, Kelleher et al., 2018, Lu and Clark, 2010]. However, the expected shift in the site frequency spectrum of positively selected cluster insertions is rather subtle and thus difficult to detect experimentally [Kofler, 2019]. In agreement with this, a recent work did not detect evidence that cluster insertions are positively selected [Zhang et al., 2020]. One drawback of this particular study is the lack of reconstruction of the entire piRNA cluster in each strain (P-element insertion sites were identified based on alignments of short reads to a reference genome) [Zhang et al., 2020]. As a consequence, P-element insertions will not be found if adjacent sequences are not conserved and the population frequency of the insertions may be estimated unreliably if the P-element inserted into repetitive regions. However, positive selection of cluster insertions could lead to an accumulation of TE insertions in piRNA clusters. Third, an insertion bias could also lead to an accumulation of TE insertions in piRNA clusters. It is likely that at least some TEs, such as the P-element, have a pronounced insertion bias into piRNA clusters [Ajioka and Eanes, 1989, Zhang et al., 2020, Kofler et al., 2018, Karpen and Spradling, 1992]. It is an important open question whether other TE families also have such an insertion bias into piRNA clusters. Alternatively, piRNA clusters may attract TE insertions, e.g. due to protein-protein interactions [Brennecke et al., 2007, Vermaak and Malik, 2009]. Finally, genetic drift could have a strong influence on the evolution of piRNA clusters. Apart from drift of cluster insertions or whole cluster haplotypes, drift may also act on the epigenetically transmitted information that determines the position of piRNA clusters. The information about the position of piRNA clusters is likely not hard coded into the DNA sequence (e.g. by motifs) but rather transmitted epigenetically by the population of maternally deposited piRNAs [Le Thomas et al., 2014a,b]. Stochastic variation in the composition and the amount of maternal transmitted piRNAs could thus lead to a rapid turnover of the location of piRNA clusters. Such a rapid turnover would likely relax selection on individual cluster insertions and make detection of positive selection on cluster insertions even more challenging.

This raises the question as to which of these processes are active in the piRNA clusters investigated in the present work. The TE content of piRNA clusters is rapidly evolving and we found that more insertions than deletions were segregating in piRNA clusters of *D. simulans*. The insertions were usually longer and occurring in younger TE families than the deletions. Most insertions are therefore likely due to recent activity of TE families in piRNA clusters. Nevertheless, some insertions (and deletions) could also be due to repeat expansion (and repeat collapse) or genomic rearrangements. A crucial question is whether the observed larger number of insertions in piRNA clusters is due to neutral processes or other forces such as positive selection on cluster insertions and an insertion bias into piRNA clusters. To distinguish between these possibilities, one would need adequate control regions, i.e. a regions that do not produce piRNAs but otherwise have very similar properties to piRNA clusters (pericentromeric regions with a similar size, number, recombination rate and TE content). It is unfortunately challenging to find suitable control regions. Additionally, larger numbers of high quality assemblies for the two *Drosophila* species may be necessary to reliably detect subtle shifts in the site-frequency spectrum of the cluster insertions as expected under positive selection. However, the properties of the deletions in piRNA clusters (short and mostly in older TEs) can likely be explained by the deletion bias observed in *Drosophila*. The gradual erosion of TEs by a deletion bias could also explain why segregating insertions (likely young) are on average longer than fixed insertions (likely old). Another important open question is whether stochastic forces or other processes such as selection and insertion biases are responsible for the differences in the rate of evolution among the piRNA clusters. It is for example possible that positive selection is stronger in clusters producing many piRNAs than in clusters producing few.

The available evidence suggests that piRNA clusters are larger in *D. melanogaster* than in *D. simulans*. This could be due to two, not mutually exclusive, reasons: first the clusters are growing in the *D. melanogaster* lineage, or second the clusters are shrinking in the *D. simulans* lineage. Our analysis of insertions and deletions suggests that even in *D. simulans* the clusters are evolving largely by insertions. If piRNA clusters were shrinking in the *D. simulans* lineage, we would not expect to see mostly insertions segregating in *D. simulans* populations. Therefore, it seems more likely that the piRNA clusters are expanding in both lineages but in *D. melanogaster* more than in *D. simulans*. This raises the question if the size of piRNA clusters could be subject to a runaway process, where larger clusters will accumulate more insertions of active TEs which, when positively selected, will lead to even larger clusters. This further raises the question whether some forces counteract the expansion of piRNA clusters. Rare and large genomic rearrangements may be an option.

We showed that the sequence and the TE content of piRNA clusters is rapidly evolving. This raises another important question - Are the positions of piRNA clusters also rapidly changing? Since the information about the position of piRNA clusters is epigenetically transmitted (see above), fluctuations in the population of maternally transmitted piRNAs could lead to changes in the size and position of piRNA clusters. This likely also happened in our investigated species. For example, the 20 investigated clusters account for 21.4% of the uniquely mapped piRNAs in *D. melanogaster* but solely for 8.4% in *D. simulans*. Hence, it is likely that other clusters, not investigated in this work, contribute the bulk of piRNAs in *D. simulans*. In agreement with this, a recent work suggests that many clusters in *Drosophila* are solely found in a single species [Gebert et al., 2021]. The turnover of the location of piRNA clusters within and among species is an important open question for future research.

Another important question is whether the observed rapid turnover of piRNA clusters is in conflict with the prevailing view on how TE invasions are stopped: the trap model holds that an invading TE is stopped when a copy of the TE jumps into a piRNA cluster [Bergman et al., 2006, Malone and Hannon, 2009, Zanni et al., 2013, Ozata et al., 2019]. For the trap model to work, it is crucial that the trap (i.e. the piRNA clusters) has a minimum size of about 0.2-3% of the genome [Kofler, 2020]. The number and genomic location of the piRNA clusters has little impact [Kofler, 2019] (except if an organism has a single piRNA cluster in non-recombining regions). As long as piRNA clusters account for at least 0.2-3% of a genome, as is likely that case in *D. melanogaster* and *D. simulans*, we do not think that the rapid turnover of piRNA clusters is in conflict with the trap model.

Finally, our work raises the question as to the consequences of rapid evolution of the composition and possibly also location of the loci responsible for silencing TEs. One consequence of such a high turnover is that silencing of TEs may be evolutionary unstable since some individuals in a population may end up without a cluster insertion for a given TE family. A high turnover of piRNA-producing loci could thus explain the low level of activity observed for many TE families in *Drosophila* [Nuzhdin, 1999] since the TE will be active in the individuals that do not produce cognate piRNAs. It is however also possible that silencing of TEs is maintained by a large number of dispersed TE insertions that are not part of piRNA cluster but nevertheless generate piRNAs [Gebert et al., 2021, Mohn et al., 2014, Shpiz et al., 2014]. These piRNA producing TEs are likely due to paramutations whereby an euchromatic TE insertion may be converted into a piRNA producing loci mediated by maternally transmitted piRNAs [Mohn et al., 2014, de Vanssay et al., 2012, Le Thomas et al., 2014b]. In agreement with this, deletion of large piRNA clusters in *D. melanogaster* did not lead to an upregulation of TEs, likely due to a large number of dispersed piRNA-producing TE insertion [Gebert et al., 2021]. If silencing against a TE is effectively based on a large and redundant number of loci, then the rapid turnover of the clusters may not lead to destabilization of the silencing of a TE, which implies that piRNA clusters may largely evolve neutrally.

## Methods

### Long-read assemblies and data

The two *D. simulans* lines *SZ232* and *SZ45* were collected in California from the Zuma Organic Orchard in Los Angeles, CA on two consecutive weekends of February 2012 [Signor et al., 2017a,b, Signor, 2020]. *SZ232* and *SZ45* were sequenced on a MinION platform (Oxford Nanopore Technologies (ONT), Oxford, GB), with fast base-calling using guppy (v4.4.2) and assembled with Canu (v2.1) [Koren et al., 2017] and two rounds of polishing with Racon (v1.4.3) and Pilon (v1.23) [Walker et al., 2014, Vaser et al., 2017, Signor et al., 2017b].

The *D. simulans* strain *m252* was collected 1998 in Madagascar and the assembly was generated with PacBio reads [Nouhaud, 2018]. The *D. simulans* strain *w*^*xD*1^ was originally collected by M. Green, likely in California, but its provenance has been lost. It is a white eyed mutant that has been maintained in the lab for more than 50 years, which can be inferred from the lack of *Wolbachia* infection [Chakraborty et al., 2021]. The *D. melanogaster* strain *A4* was sampled 1963 in Koriba Dam (Zimbabwe) [King et al., 2012]. The reference strain *Iso-1* of *D. melanogaster* was generated by crossing several laboratory strains, with largely unknown sampling data [Brizuela et al., 1994]. *Canton-S* was sampled 1935 in Ohio (USA) [Anxolabéhè re et al., 1988]. We could not obtain details on the sampling of the *D. sechellia* strain *sech25* (*Robertson 3C*) and the *D. mauritiana* strain *mau12* (*w12*) [Chakraborty et al., 2021]. The assemblies of the *D. melanogaster* strain *A4* (ASM340174v1), the *D. simulans* strain *w*^*xD*1^ (ASM438218v1), the *D. sechellia* strain *sech*25 (ASM438219v1) and the *D. mauritiana* strain *mau*12 (ASM438214v1) are based on PacBio reads [Chakraborty et al., 2018, 2021]. The assembly of the *D. melanogaster* strain *Canton-S* was generated using ONT reads [Wierzbicki et al., 2021]. We obtained the assembly of the *D. melanogaster* reference strain *Iso-1* from FlyBase (r6; [Hoskins et al., 2015].

### Identifying homologous piRNA clusters

Previously, we designed flanking sequences for 85 out of the 142 annotated piRNA clusters in *D. melanogaster* [Wierzbicki et al., 2021]. We excluded piRNA clusters at the end of chromosomes where two flanking sequences cannot be found, as well as clusters on the fragmented U chromosome. The *D. melanogaster* flanking sequences were aligned to each assembly using bwa bwasw (0.7.17-r1188; [Li and Durbin, 2010]). The alignments were repeated using bwa mem -a (to show alternative hits) to identify clusters that were not recovered by bwa bwasw. Homologous clusters were identified as the regions between the aligned *D. melanogaster* flanking sequences [Wierzbicki et al., 2021]. Cluster sequences with internal gaps were excluded. We validated the homology of clusters with a reciprocal mapping approach. First, we designed independent sets of flanking sequences in the target strain (e.g. *D. simulans*) that did not overlap with the aligned *D. melanogaster* flanking sequences. Second we aligned these reciprocal flanking sequences with bwa bwasw and bwa mem -a to release 5 of the *D. melanogaster* reference genome (piRNA clusters were annotated in release 5 [Brennecke et al., 2007]). Finally, we checked whether the coordinates of the annotated piRNA clusters were contained within the positions of the aligned reciprocal flanking sequences (supplementary tables S1-S3).

### Assembly quality of piRNA clusters

Even when both flanking sequences align to the same contig, a piRNA cluster may be incorrectly assembled, for example if some internal sequences are missing in the assembly. We previously proposed that heterogeneity of the base coverage (e.g. due to repeat collapse) and an elevated soft-clip coverage (resulting from unaligned read termini) can be used to identify assembly errors in clusters [Wierzbicki et al., 2021]. To examine these patterns in our assemblies, we aligned the long reads used for generating the assembly back to the respective assembly using minimap2 (v2.16-r922; v2.17-r954) [Li, 2018]. The exception to this was *D. melanogaster Iso-1* where the long reads are not from the original assembly but from a slightly diverged sub-strain Solares et al. [2018]. As reference, we computed the 99% quantiles of the base and soft-clip coverage of complete BUSCO (Benchmarking Universal Single-Copy Orthologs (v3.0.2; v5.0.0); [Simão et al., 2015]) genes based on the *Diptera odb9* or *Diptera odb10* data set. Regions where the base or the soft-clip coverage markedly deviates from the 99% quantile of the BUSCO genes could indicate an assembly error and serve as a guide to the quality of the overall cluster assembly.

### Aligning the annotations of piRNA clusters

To align the TE annotations of homologous piRNA clusters, we first extracted the sequences of the clusters from the assemblies with samtools (v1.9; [Li et al., 2009]) based on the positions of the aligned flanking sequences. Next, we annotated TEs in these sequences using RepeatMasker (open-4.0.7) with a *D. melanogaster* TE library and the parameters: -s (sensitive search), -nolow (disable masking of low complexity sequences), and -no is (skip check for bacterial IS) [Smit et al., 2013-2015, Bao et al., 2015, Quesneville et al., 2005]. Finally, we aligned the resulting repeat annotations with our novel tool Manna (see Results) using the parameters -gap 0.09 (gap penalty), -mm 0.1 (mismatch penalty) -match 0.2 (match score).

### Visualising piRNA clusters

For visualizing the composition and evolution of piRNA clusters, we annotated repeats in piRNA clusters using the *D. melanogaster* TE library and RepeatMasker (open-4.0.7; [Smit et al., 2013-2015, Bao et al., 2015, Quesneville et al., 2005]. Homologous sequences in piRNA clusters were identified with blastn (BLAST 2.7.1+ [Altschul et al., 1990]) using default parameters. We visualized the annotation and the sequence similarity of piRNA clusters with Easyfig (v2.2.3 08.11.2016) [Sullivan et al., 2011] setting the similarity scale to a minimum of 70%. Finally, we merged the pairwise visualizations generated by Easyfig to allow comparing multiple clusters. A walkthrough for this pipeline is available at https://sourceforge.net/p/manna/wiki/piRNAclusterComparison-walkthrough/.

### piRNAs

We obtained previously published piRNA data from ovaries of *D. simulans* (ERR1821669) and *D. melanogaster* (ERR1821654) strains sampled from Chantemesle (France) [Asif-Laidin et al., 2017]. We trimmed the adaptor sequence (TGGAATTCTCGGGTGCCAAG) with cutadapt (v3.4; [Martin, 2011]). The reads were aligned to the reference genomes (*D. melanogaster* : *Iso-1, D. simulans*: *w*^*xD*1^ with novoalign (V3.03.02; http://novocraft.com/). The coordinates of the piRNA clusters were obtained from the aligned flanking sequences (see above). We retained reads with a length between 23 and 29bp, normalized the abundance of these reads to a million mapped reads and visualized the abundance of ambiguously (*mq* = 0) and unambiguously (*mq >* 0) mapped reads along piRNA clusters with R (v3.6.1) and ggplot2 (v3.3.3)[R Core Team, 2012, Wickham, 2016].

## Availability

The reads and the assemblies of the two *D. simulans* strains are publicly available (PRJNA736739; PRJNA736415). The novel software for a multiple alignments of annotations, Manna, is available at https://sourceforge.net/projects/manna/. A manual and the validations are available at https://sourceforge.net/p/manna/wiki/Home/. The TE library and list of TE names used in this work are available at https://sourceforge.net/projects/manna/files/pirnaclustercomparison/resources/. All script used in this work are available at https://sourceforge.net/projects/manna/files/publicationdata/

## Author contributions

FW, RK, and SS conceived this work. SS assembled the two *D. simulans* strains. RK developed Manna. FW, RK and SS analyzed the data. FW, RK and SS wrote the manuscript.

## Acknowledgments

We thank all members of the Institute of Population Genetics for feedback and support. This work was supported by the Austrian Science Foundation (FWF) grant P30036-B25 to RK and by the National Science Foundation Established Program to Stimulate Competitive Research (NSF-EPSCoR-1826834), the North Dakota EPSCoR STEM grants program, and NSF-EPSCoR-2032756 to SS.

