## Supplementary Material for "Evolutionary dynamics of piRNA clusters in *Drosophila*"

### Supplementary figures and tables

August 20, 2021

#### Supplementary figures

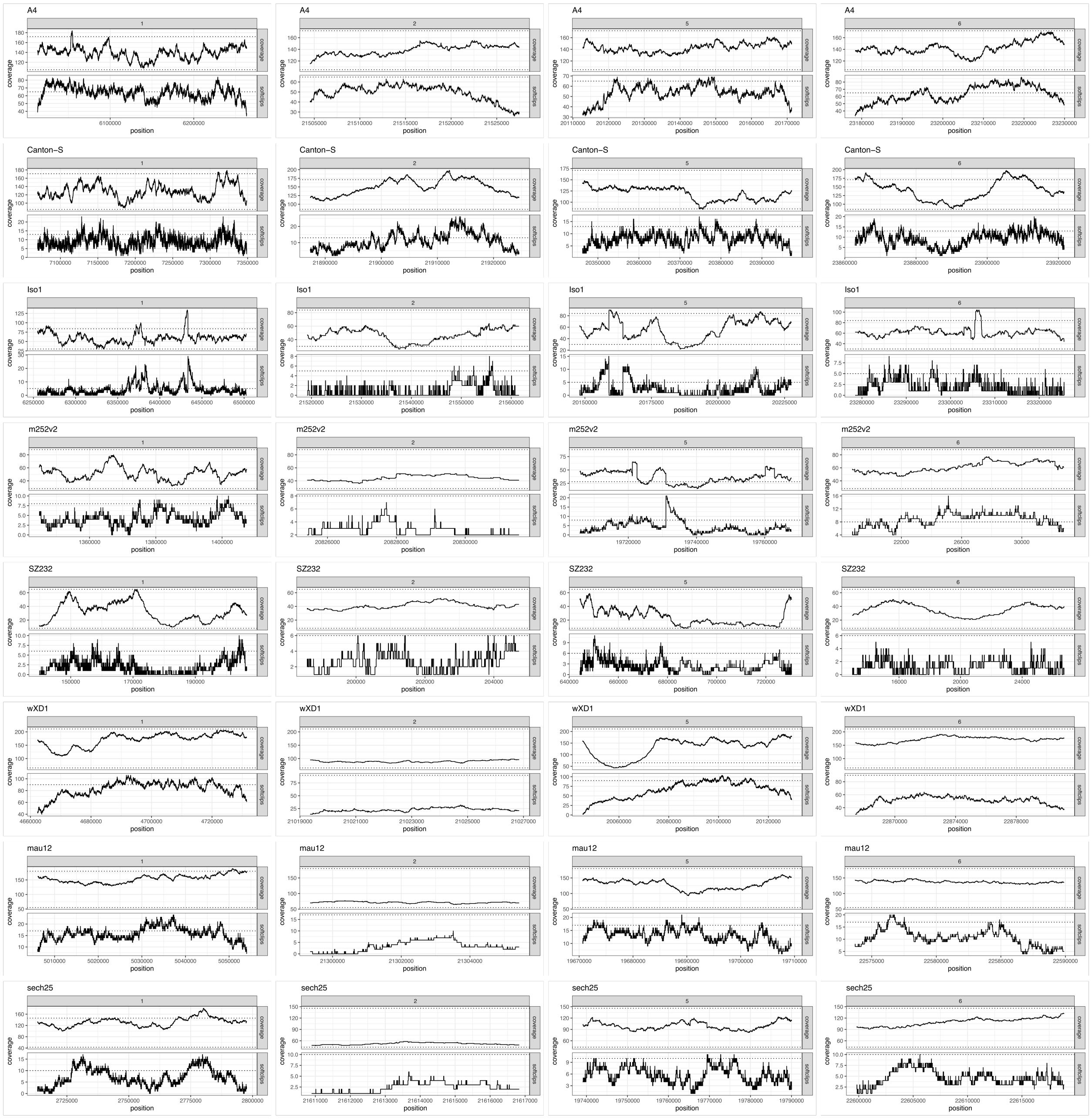

Figure S1: Quality metrics for clusters 1, 2, 5 and 6. Each plot shows the base and the softclip coverage based on an alignment of long-reads. Dotted lines show the 99% quantiles for complete BUSCO genes. There could be an assembly error in cluster 5 of strain m252v2 and clusters 1 and 5 of strain Iso-1. We note that the long-reads used for computing the quality metrics in Iso-1 were not used in the assembly of the strain and may thus be derived from a slightly diverged sub-strain [Rahman et al., 2015].

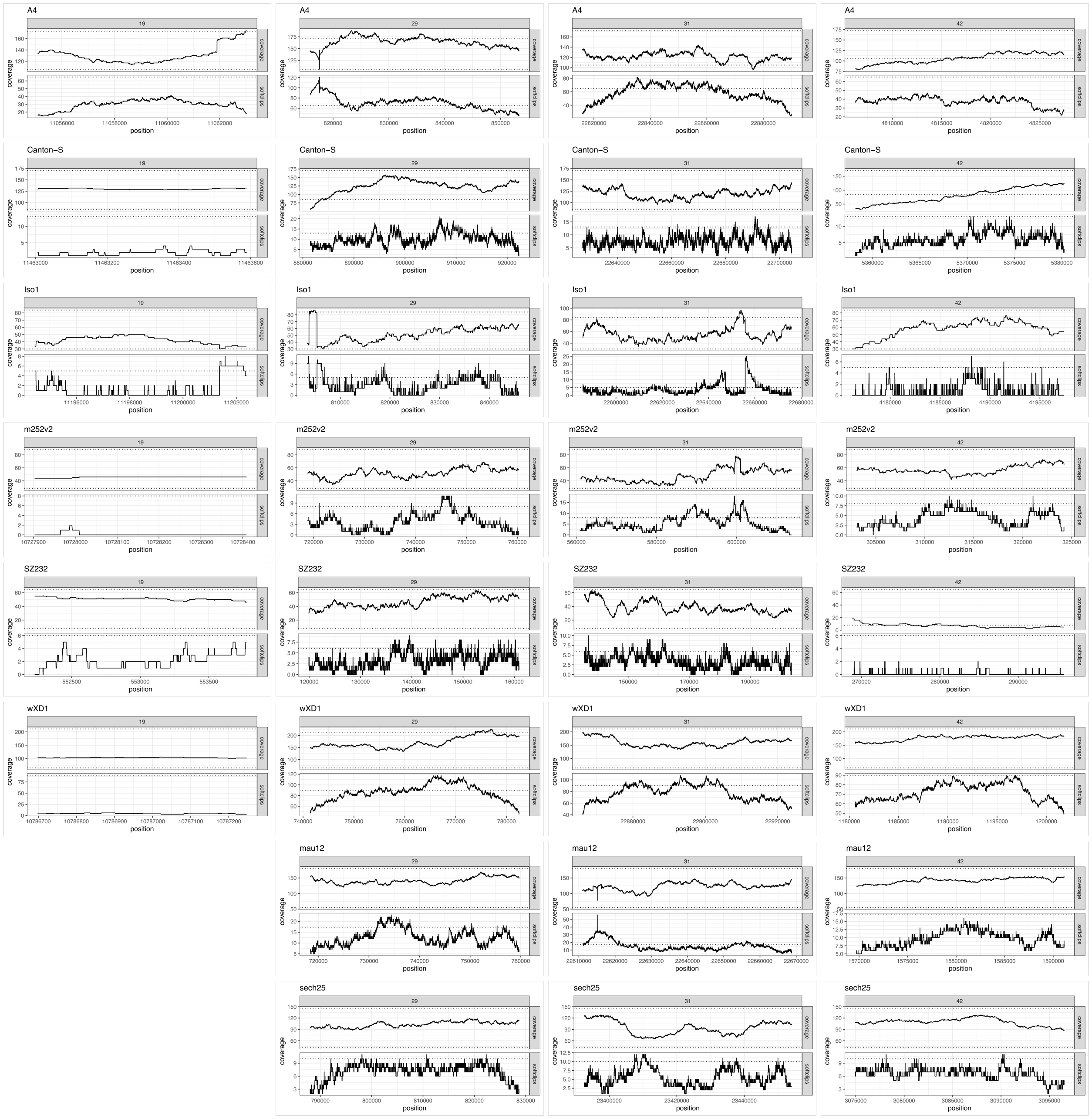


Figure S2: Quality metrics for clusters 19, 29, 31 and 42. Each plot shows the base and the softclip coverage based on an alignment of long-reads. Dotted lines show the 99% quantiles for complete BUSCO genes. Cluster 19 was empty in the strains mau12 and sech25. There could be an assembly error in cluster 29 of strains A4 and Iso-1 and cluster 31 of strains mau12 and Iso-1. We note that the long-reads used for computing the quality metrics in Iso-1 were not used in the assembly of the strain and may thus be derived from a slightly diverged sub-strain [Rahman et al., 2015].

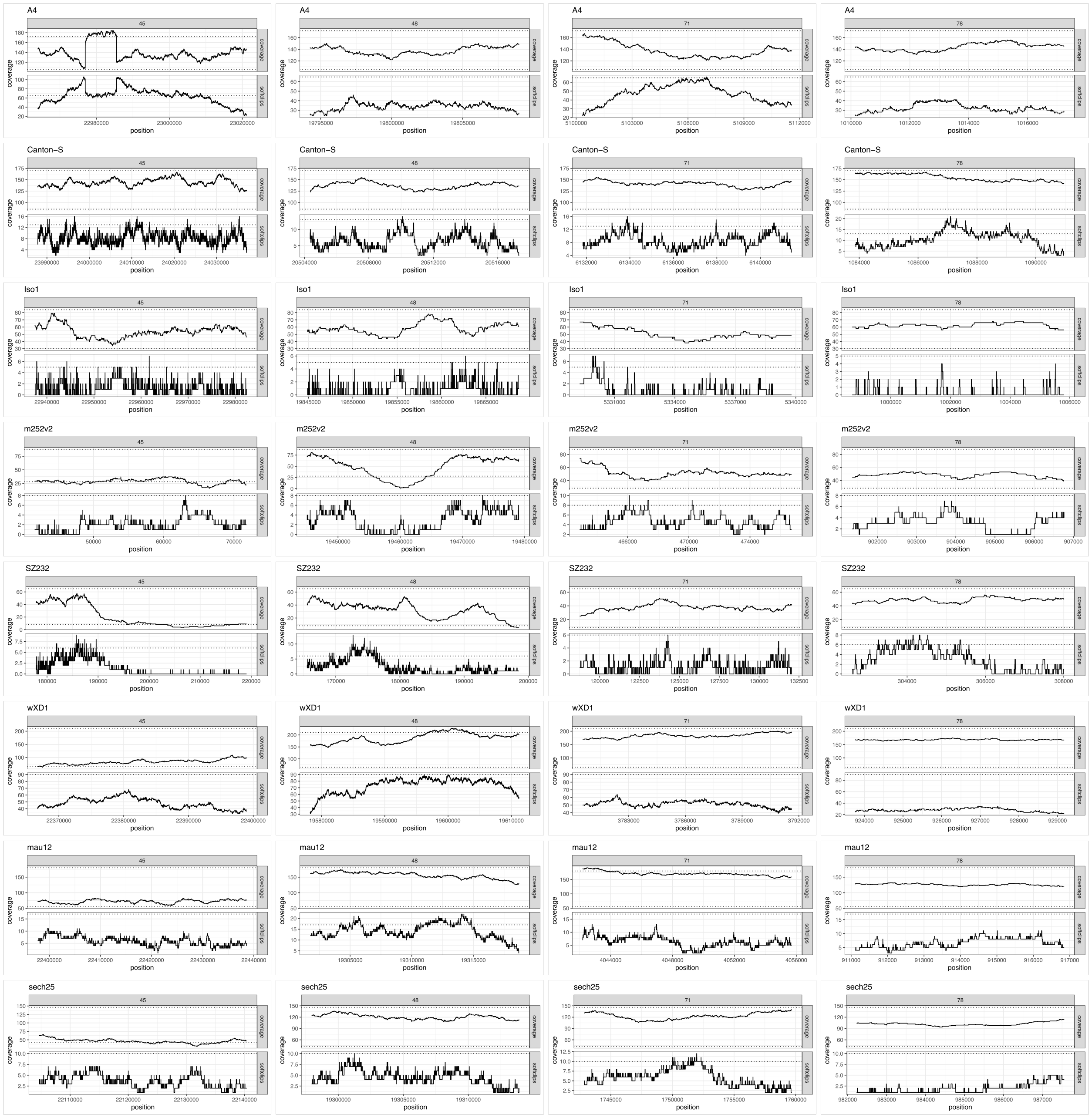


Figure S3: Quality metrics for clusters 45, 48, 71 and 78. Each plot shows the base and the softclip coverage based on an alignment of long-reads. Dotted lines show the 99% quantiles for complete BUSCO genes. There could be an assembly error in cluster 45 of stain A4 .

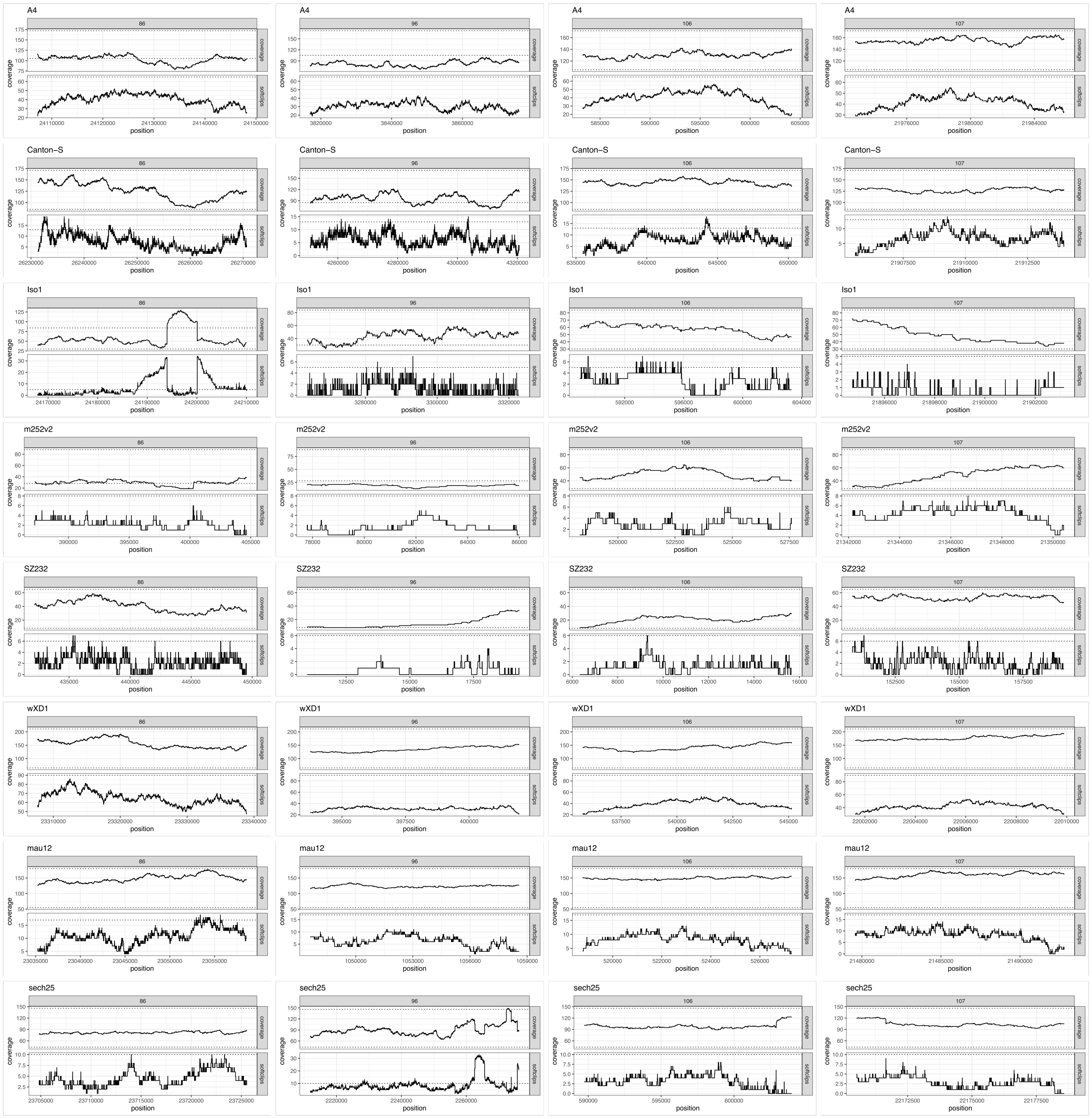


Figure S4: Quality metrics for cluster 86, 96, 106 and 107. Each plot shows the base and the softclip coverage based on an alignment of long-reads. Dotted lines show the 99% quantiles for complete BUSCO genes. There could be an assembly error in cluster 96 of strain sech25 and cluster 86 of strain Iso-1. We note that the long-reads used for computing the quality metrics in Iso-1 were not used in the assembly of the strain and may thus be derived from a slightly diverged sub-strain [Rahman et al., 2015].

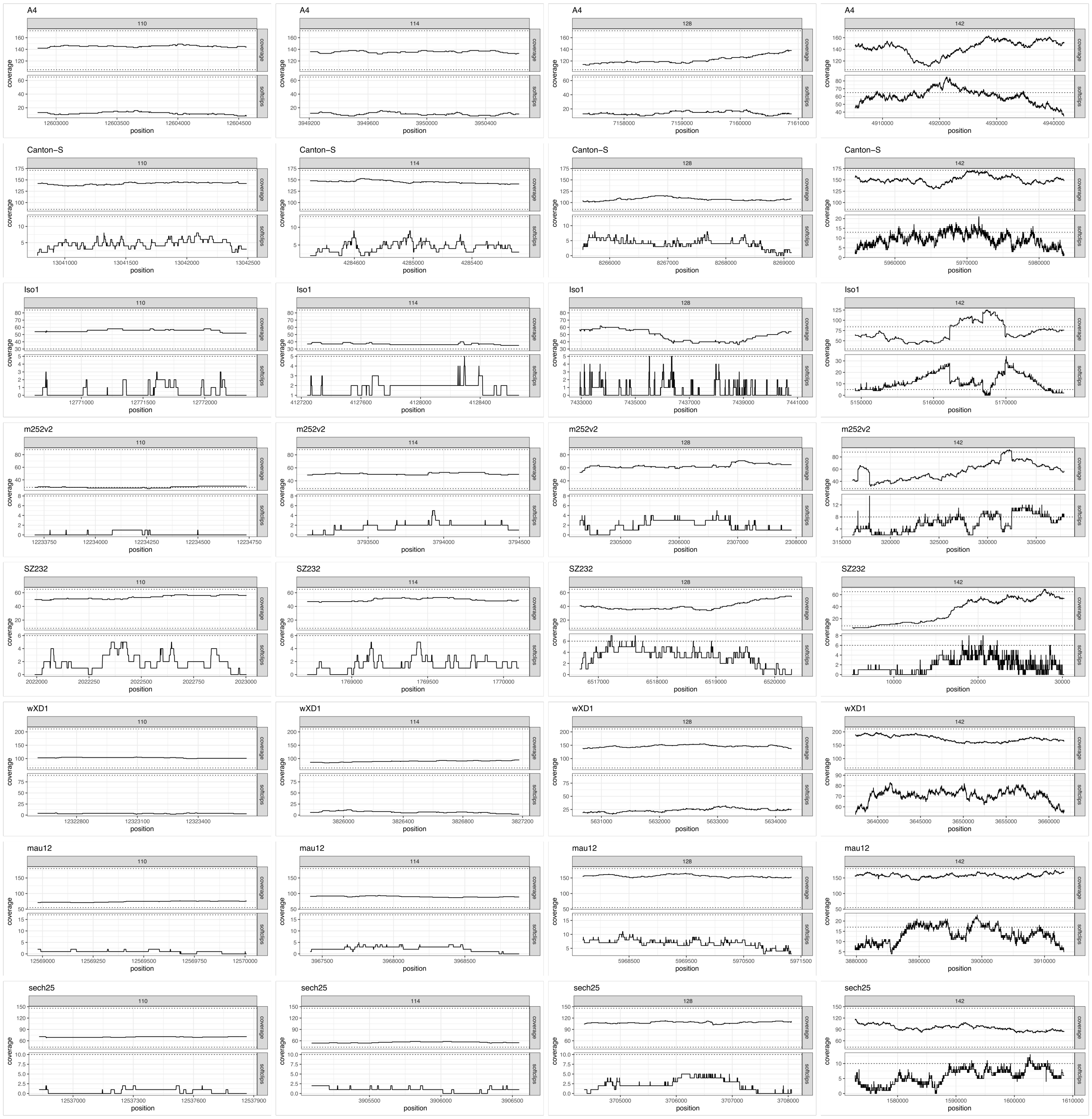


Figure S5: Quality metrics for cluster 110, 114, 128 and 142. Each plot shows the base and the softclip coverage based on an alignment of long-reads. Dotted lines show the 99% quantiles for complete BUSCO genes. There could be an assembly error in cluster 142 of A4, Iso-1 and m252v2. We note that the long-reads used for computing the quality metrics in Iso-1 were not used in the assembly of the strain and may thus be derived from a slightly diverged sub-strain [Rahman et al., 2015].

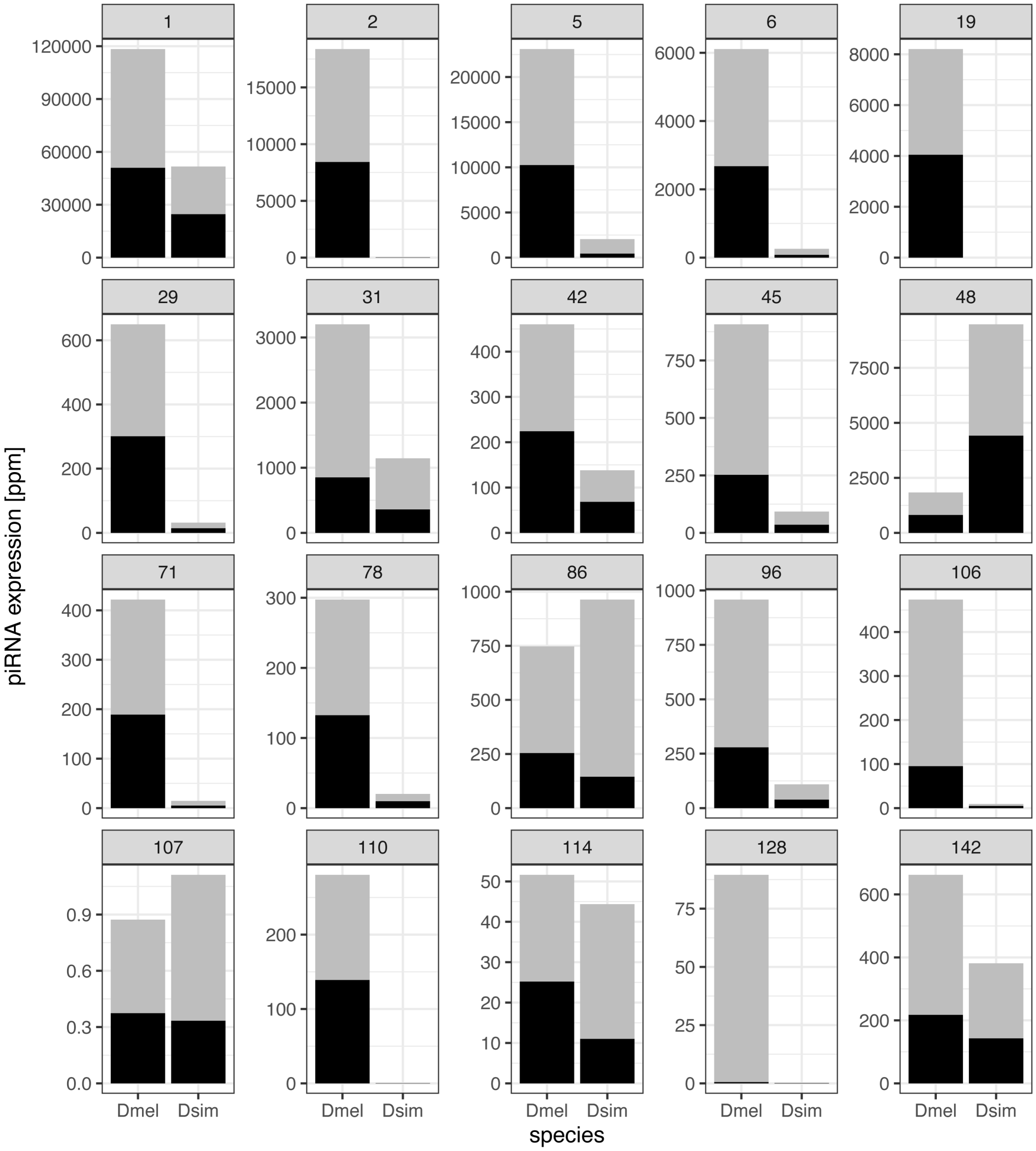


Figure S6: Expression of piRNAs in homologous piRNA clusters of *D. simulans* and *D. melanogaster*. The abundance of piRNAs was normalized to a million mapped reads with a length between 23-29nt. Unambiguously mapped piRNAs ($mq > 0$) are shown in dark grey whereas ambiguously mapped piRNAs ($mq = 0$) are in light grey.

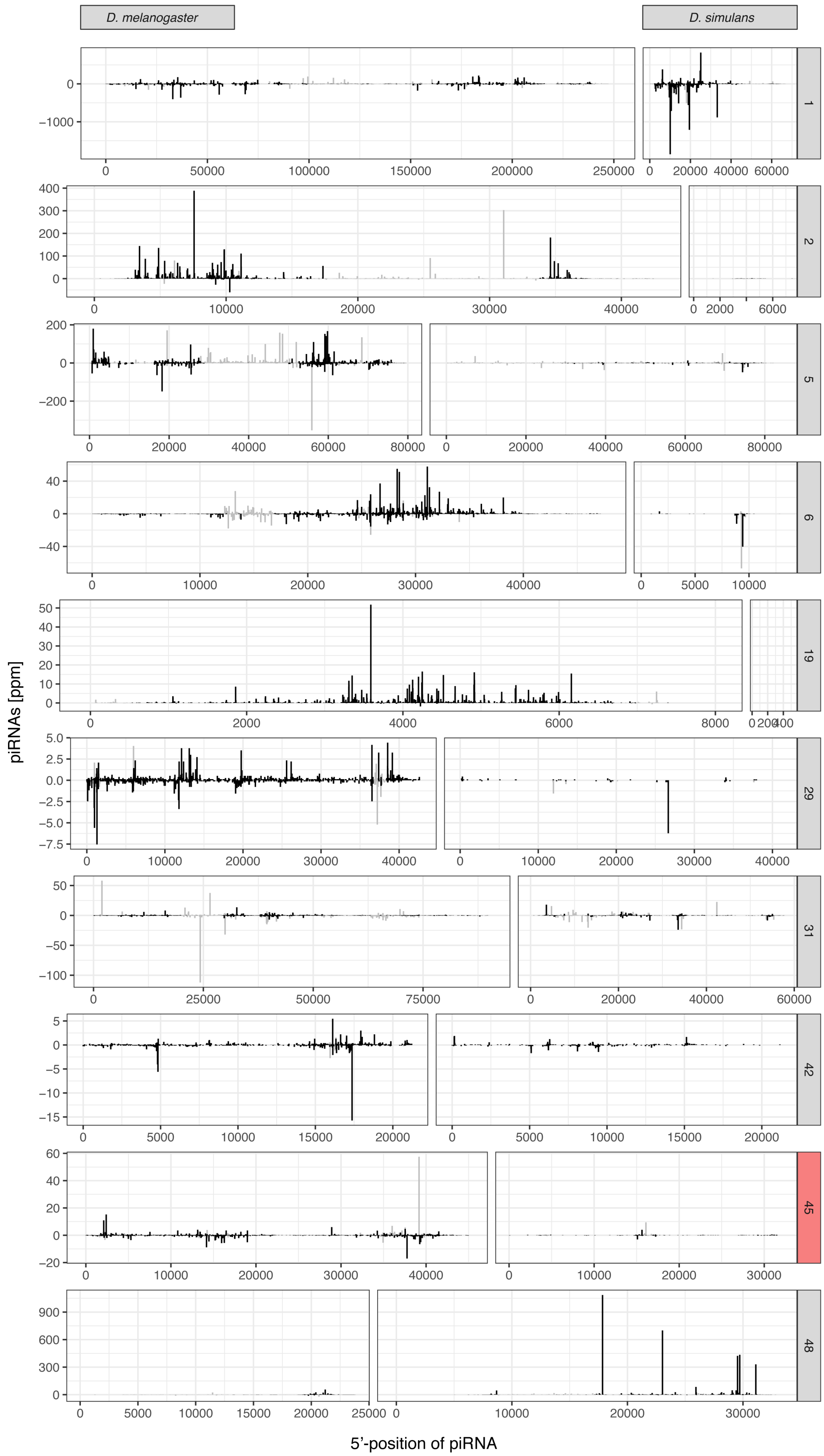


Figure S7: Distribution of piRNAs along homologous piRNA clusters of *D. simulans* and *D. melanogaster*. Sense piRNAs are shown on the positive y-axis and antisense piRNAs on the negative y-axis. Unambiguously mapped piRNAs ($mq > 0$) are in dark grey whereas ambiguously mapped piRNAs ($mq = 0$) are in light grey. Except for cluster 45 which is reverse complemented in *D. simulans*, all clusters are shown in the same 5'-to-3' orientation in the two species.

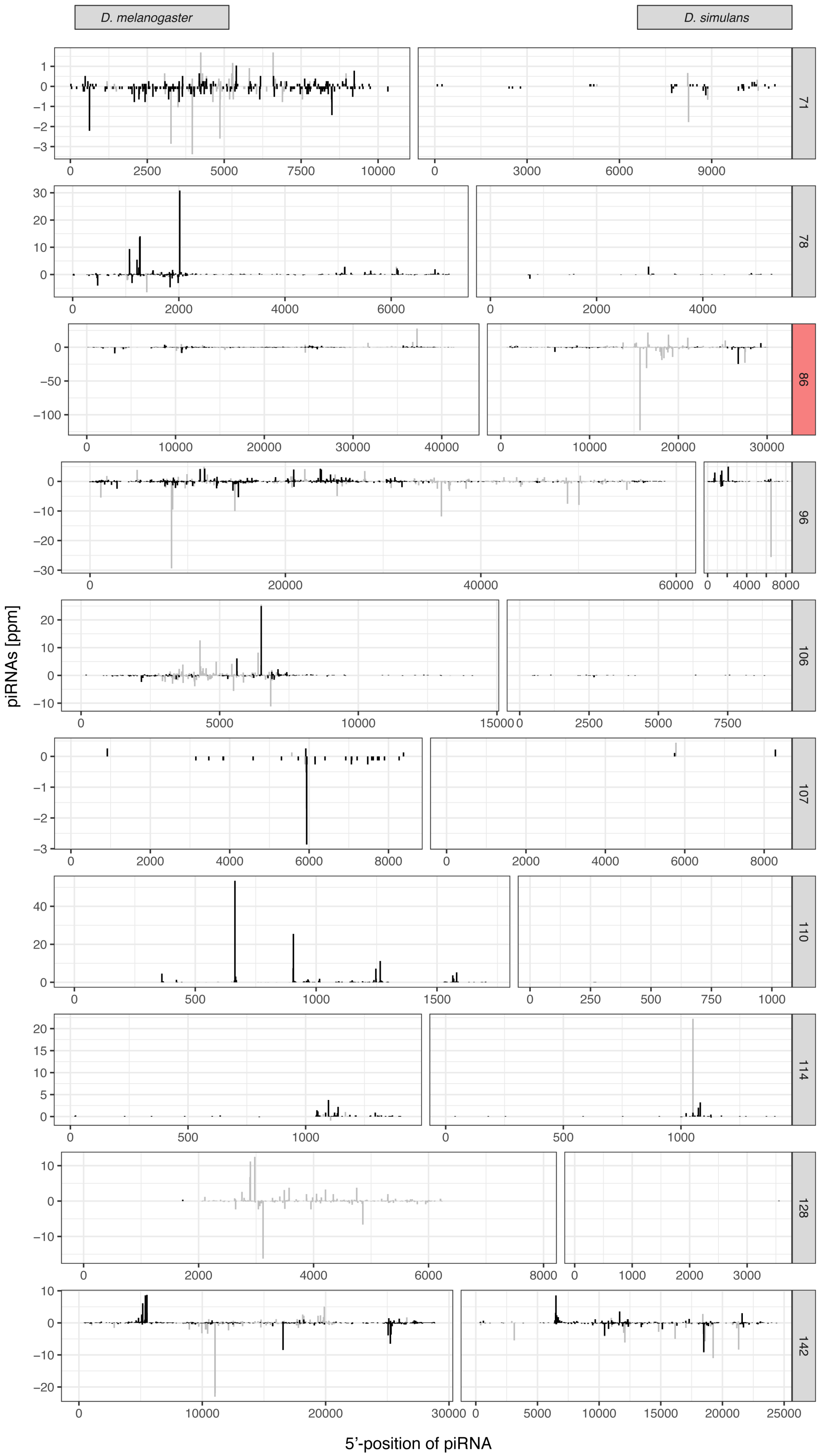


Figure S8: Distribution of piRNAs along homologous piRNA clusters of *D. simulans* and *D. melanogaster*. Sense piRNAs are shown on the positive y-axis and antisense piRNAs on the negative y-axis. Unambiguously mapped piRNAs ($mq > 0$) are in dark grey whereas ambiguously mapped piRNAs ($mq = 0$) are in light grey. Except for cluster 86 which is reverse complemented in *D. simulans*, all clusters are shown in the same 5'-to-3' orientation in the two species.

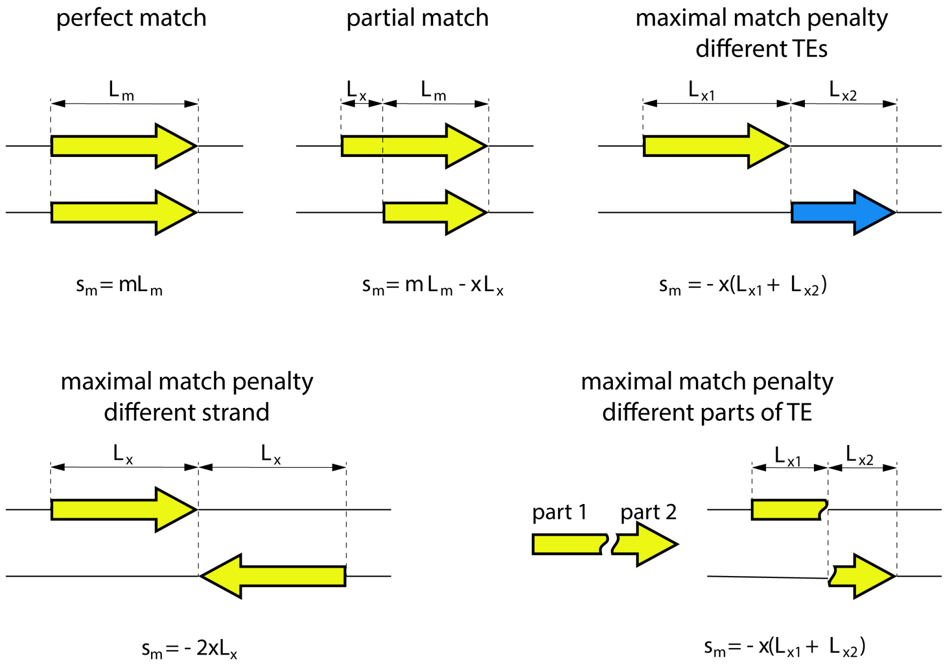


Figure S9: Examples illustrating computation of the match score ($s_m$) in Manna. The match score is computed based on the length of the sequence matching among two TE insertions ($L_m$), the length of the not matching sequence ($L_x$) the match score (m) and the mismatch score ($x$). As usual for Needeleman-Wunsch algorithm, the score for each position in the similarity matrix is computed as $S = max(s_m, s_i, s_d)$ where $s_i = L_i * g$ is the gap score of an insertion and $s_d = L_d * g$ the gap score of a deletion. $g$ gap score, $L_i$ length of inserted sequence; $L_d$ length of deleted sequence

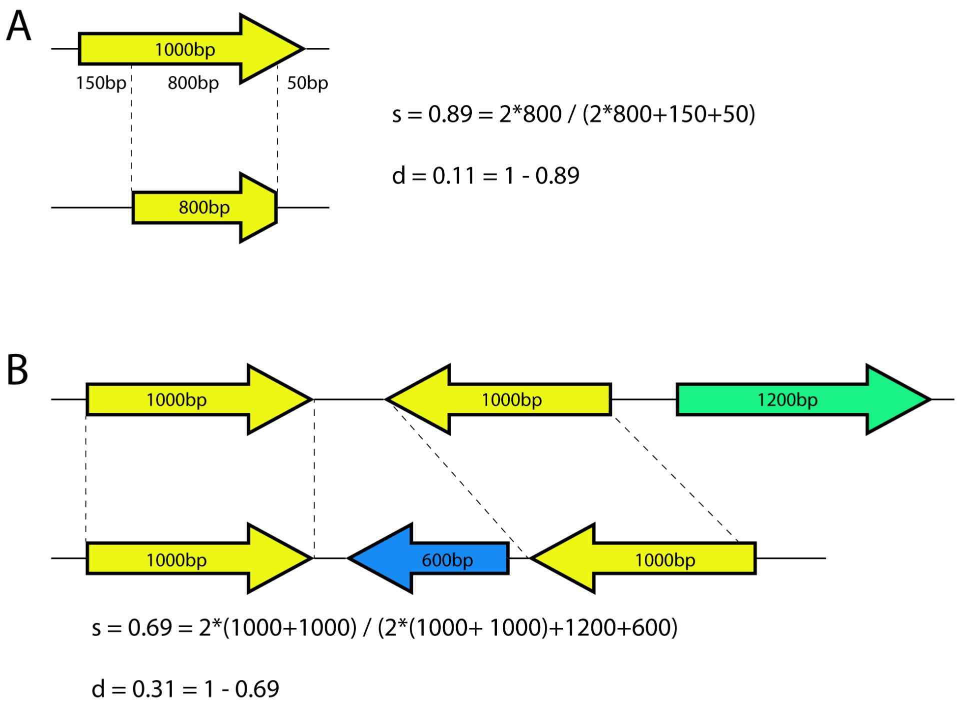


Figure S10: Examples on how we computed the similarity ($s$) and distance ($d$) among two TE annotations aligned with Manna. The similarity was computed as $s = 2 * a/2 * a + u$ where $a$ and $u$ is the total length of the aligned and unaligned TEs. The distance is computed as $d = 1 - s$. TE insertions are shown as colored arrows, where the direction of the arrow indicates the strand and the color the family of the insertion. A) Example for an alignment where one TE has two terminal deletions. Aligned sequences are marked by dashed lines B) Complex example involving several aligned and unaligned TE sequences

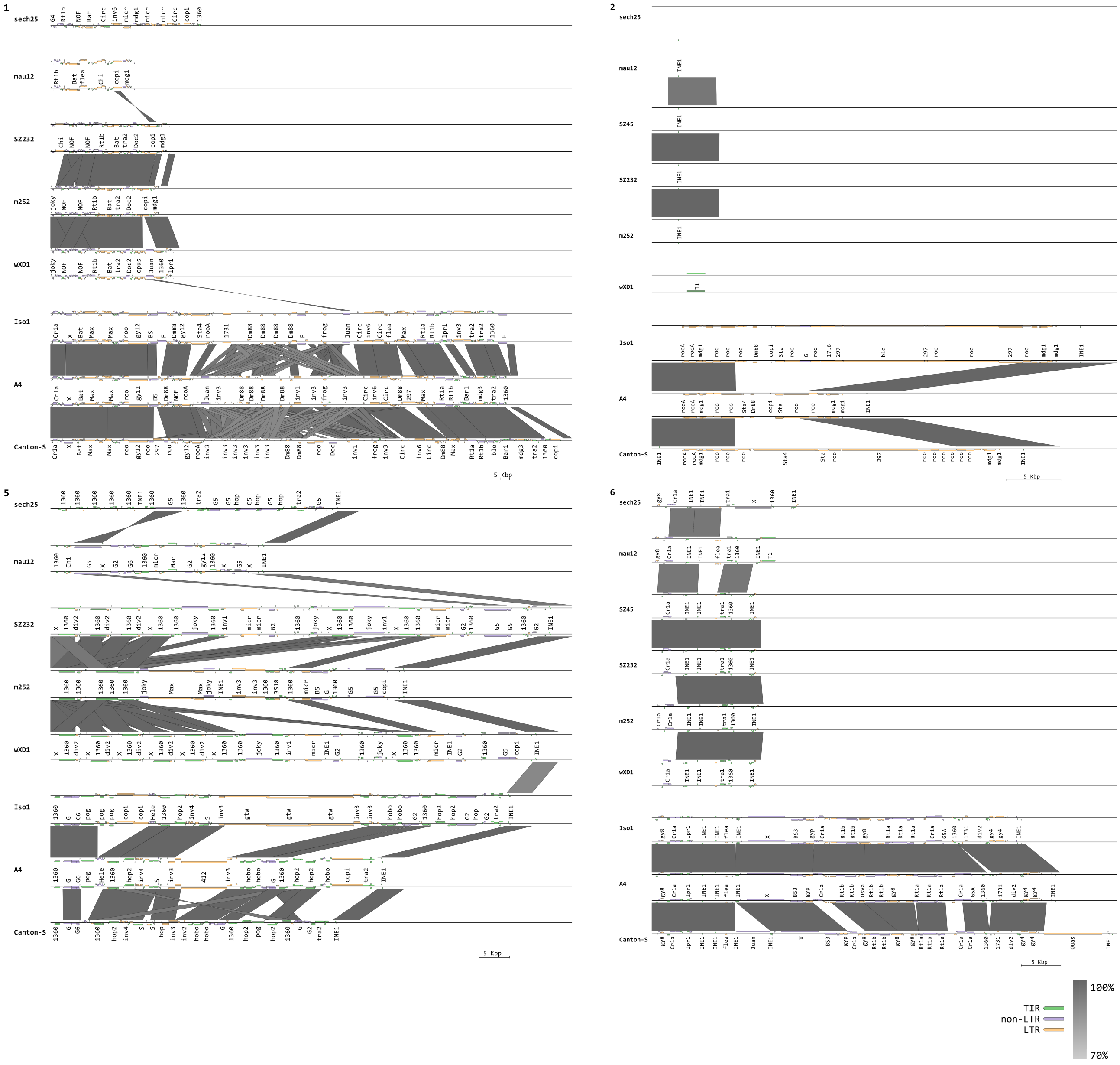


Figure S11: Composition of piRNA clusters 1, 2, 5 and 6 in different Drosophila species. Grey bars indicate regions of similarity among two assemblies (minimum length 3 kb). TE families are colored by order (LTR, non-LTR and TIR).

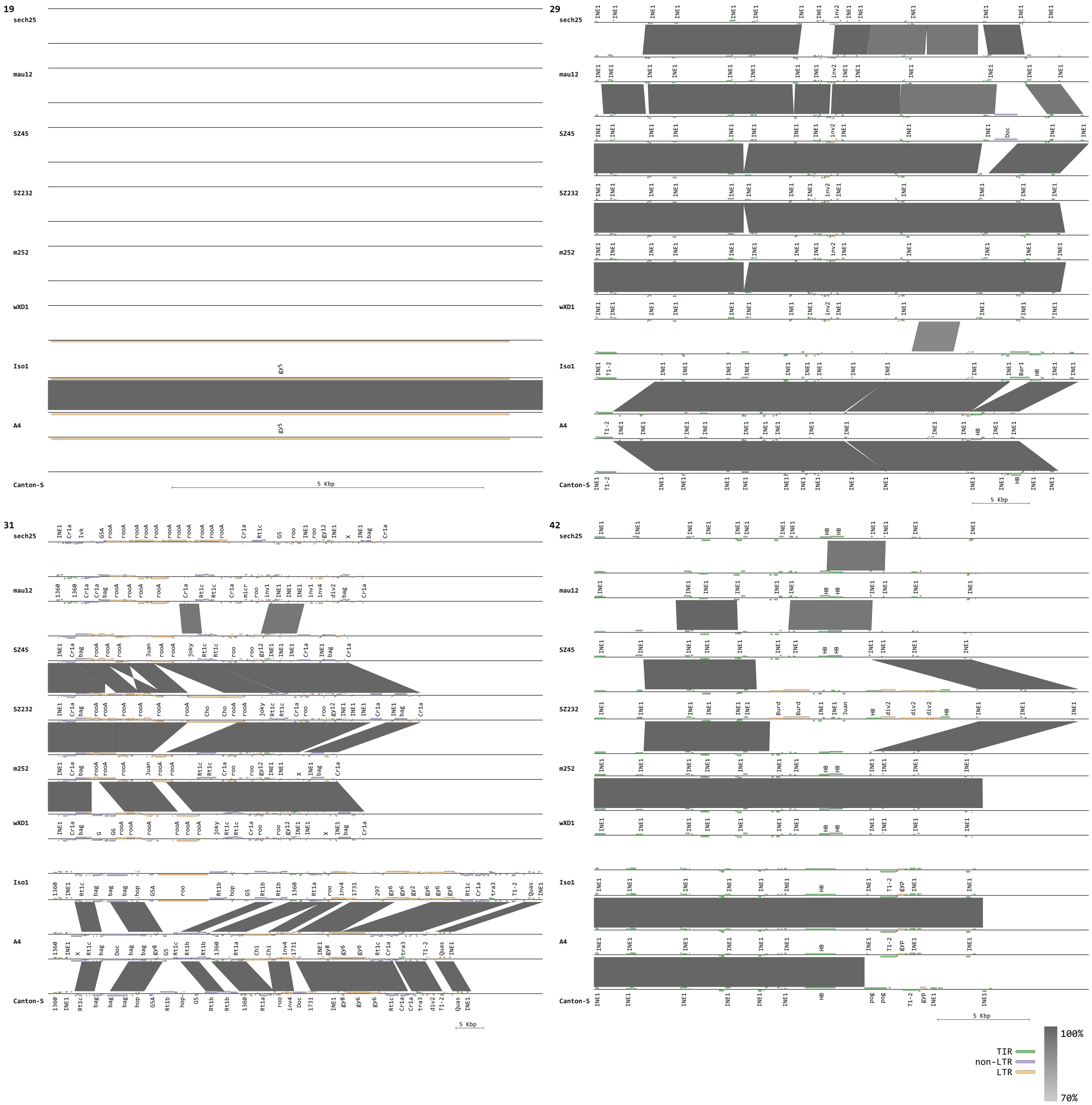

Figure S12: Composition of piRNA clusters 19, 29, 31 and 42 in the different Drosophila species. Grey bars indicate regions of similarity among two assemblies (minimum length 3 kb). TE families are colored by order (LTR, non-LTR and TIR).

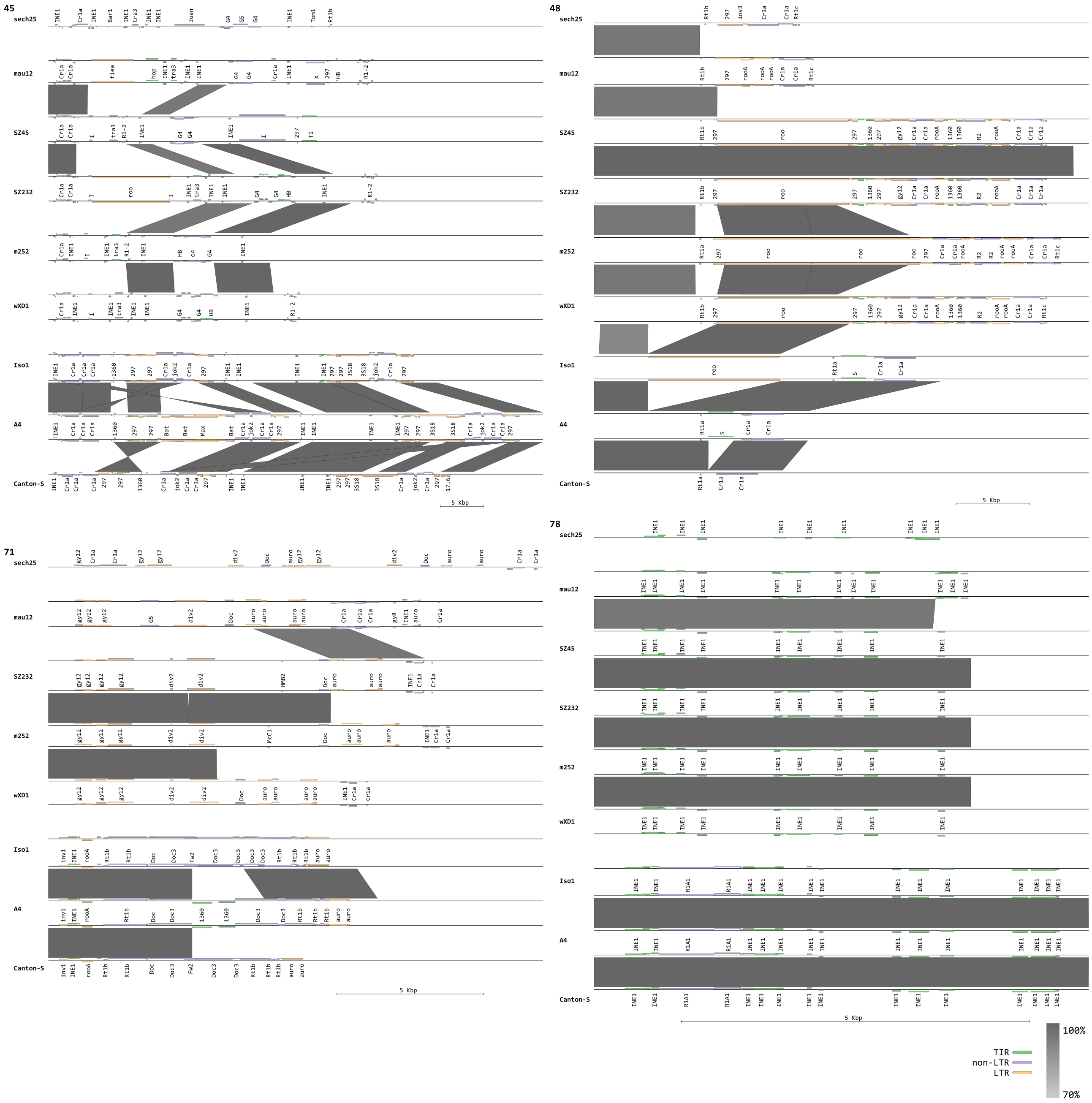


Figure S13: Composition of piRNA clusters 45, 48, 71 and 78 in the different Drosophila species. Grey bars indicate regions of similarity among two assemblies (minimum length 3 kb). TE families are colored by order (LTR, non-LTR and TIR).

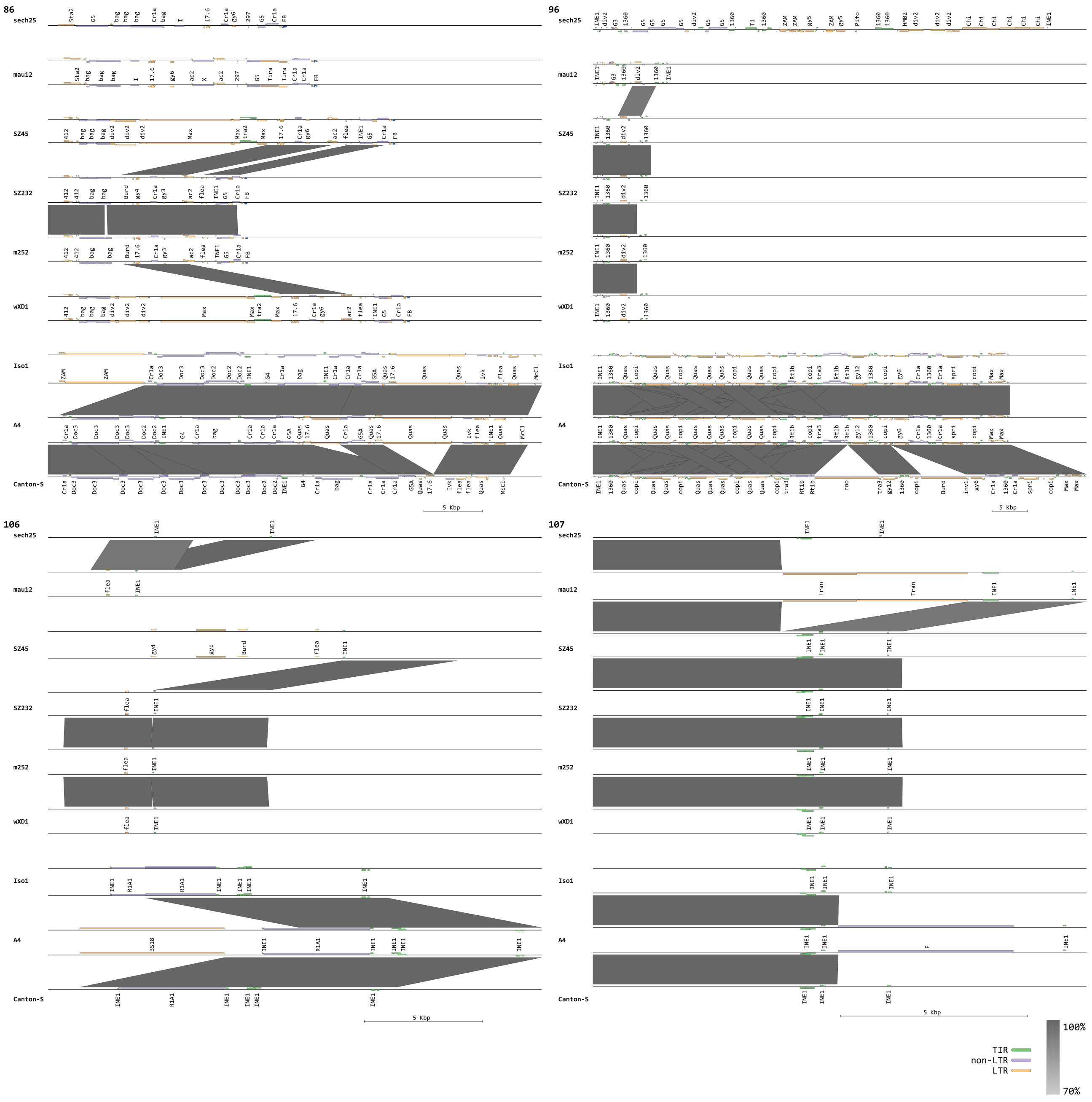


Figure S14: Composition of piRNA clusters 86, 96, 106 and 107 in the different Drosophila species. Grey bars indicate regions of similarity among two assemblies (minimum length 3 kb). TE families are colored by order (LTR, non-LTR and TIR).

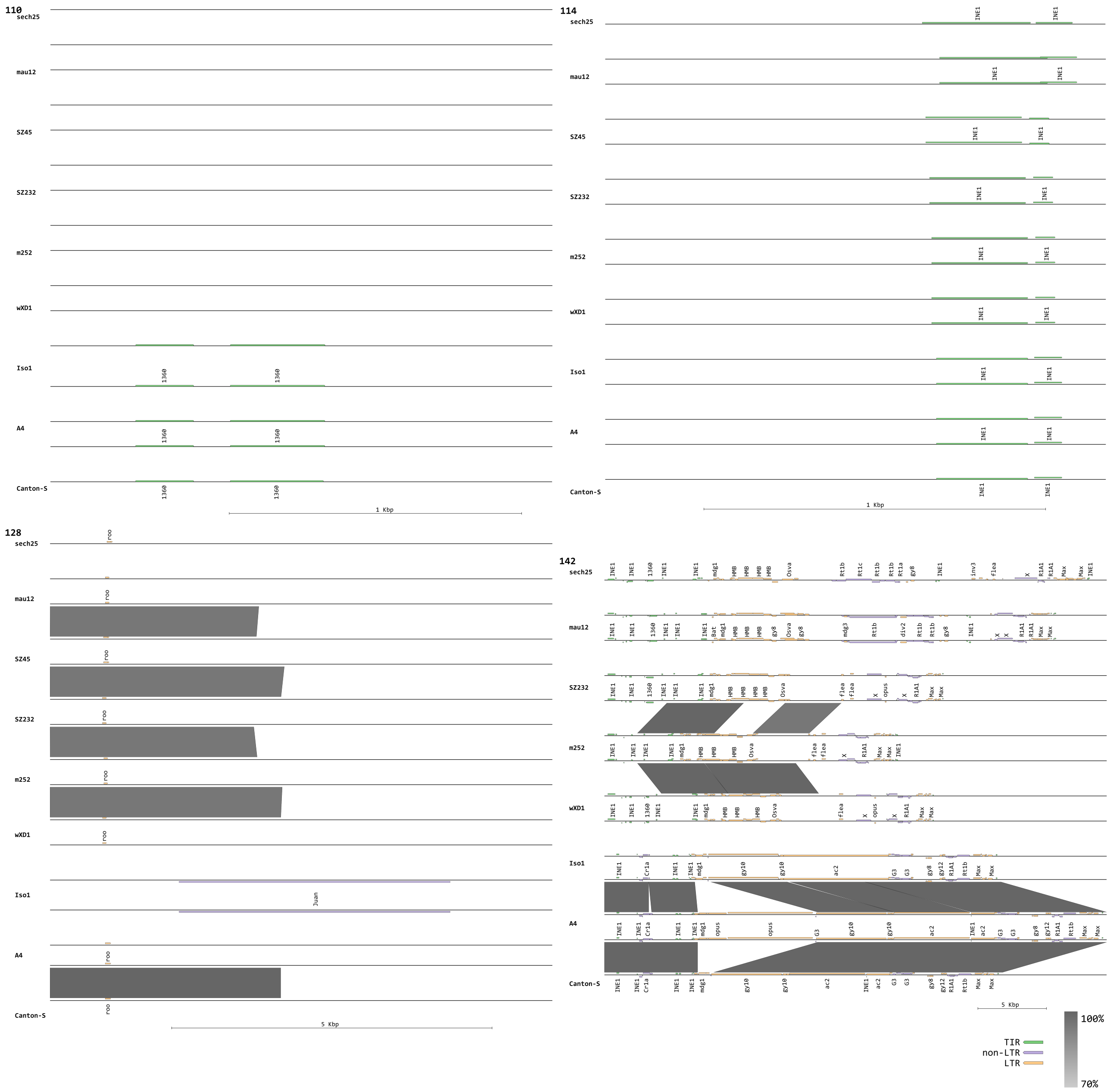


Figure S15: Composition of piRNA clusters 110, 114, 128 and 142 in the different Drosophila species. Grey bars indicate regions of similarity among two assemblies (minimum length 3 kb). TE families are colored by order (LTR, non-LTR and TIR).

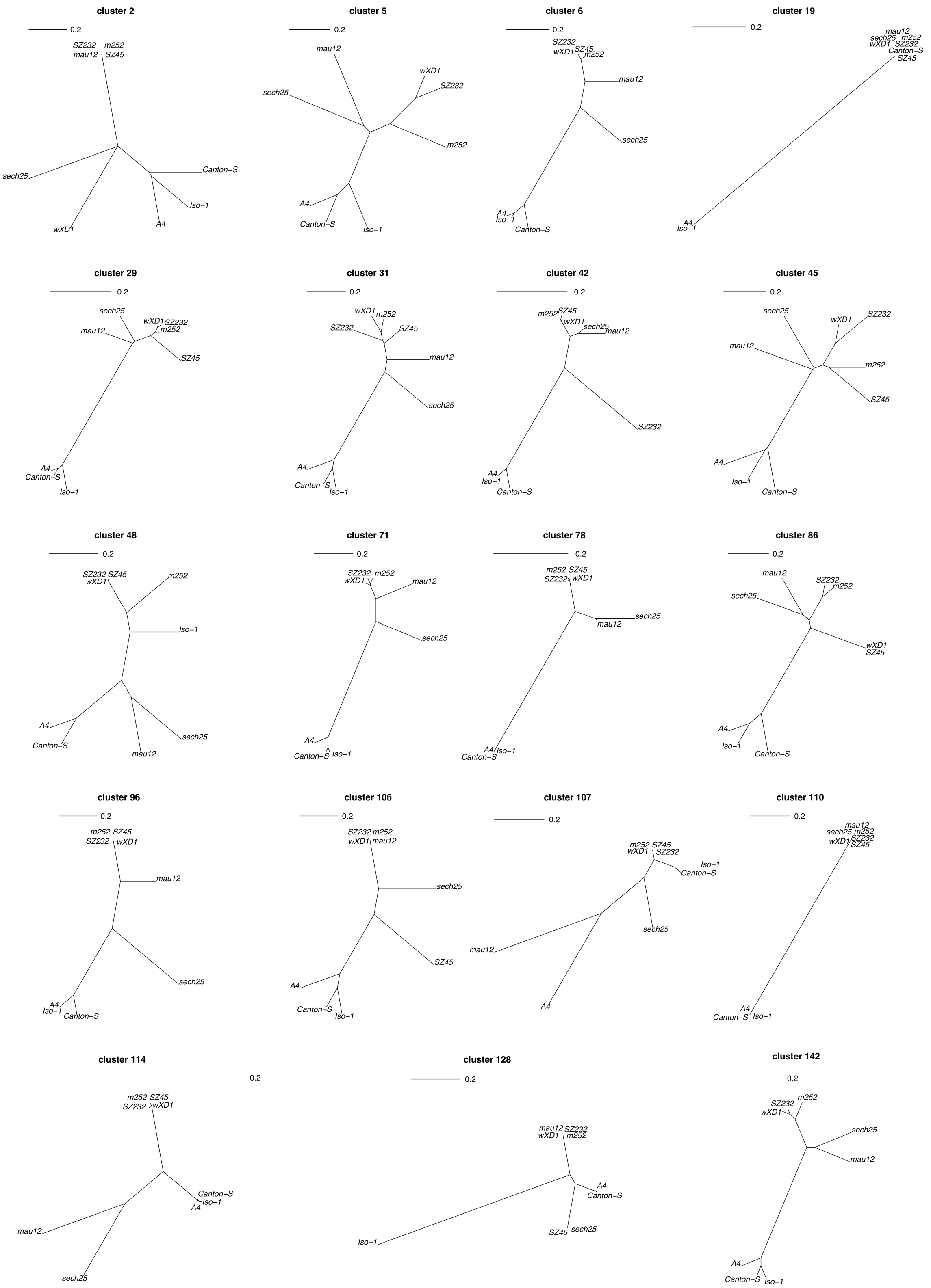


Figure S16: Phylogenetic trees showing the distance among the strains/species for 19 clusters. The tree for cluster 42AB (cluster 1) is shown in the main manuscript. The evolutionary distance is estimated by Manna as the fraction of unaligned sequences. The scale bars show a distance of 0.2 (i.e. 20% unaligned TE sequences). Solely 16 cluster were completely assembled for SZ45. *D. simulans*: SZ45, SZ232, m252, $w^{xD1}$; *D. melanogaster*: A4, Canton-S, Iso-1; *D. mauritiana*: mau12; *D. sechellia*: sech25

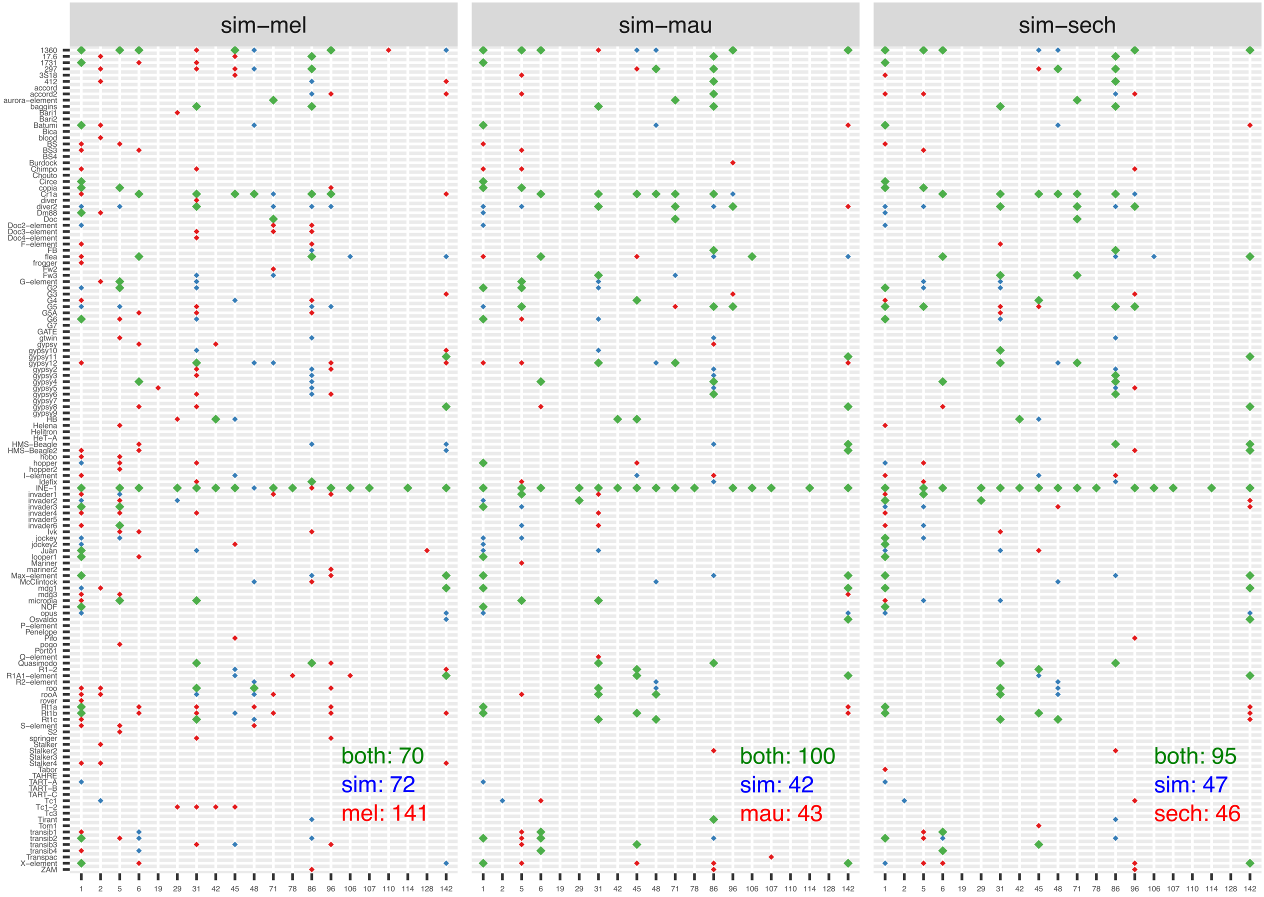


Figure S17: Overview of the TE content of piRNA clusters in *D. simulans* (sim), *D. melanogaster* (mel), *D. mauritiana* (mau) and *D. sechellia* (sech). For each piRNA cluster (x-axis) we indicate whether a given TE family (y-axis) has at least one insertion in *D. simulans* (blue), the other species (red; dependent on the comparison either mel, mau or sech) or both species (green). Summary counts are shown in corresponding plots. For each species solely the content of a single strain was anaylsed (*Iso-1* for *D. melanogaster* and $w^{xD1}$ for *D. simulans*; in contrast to the main figure which is based on three strains for *D. melanogaster* and *D. simulans*).

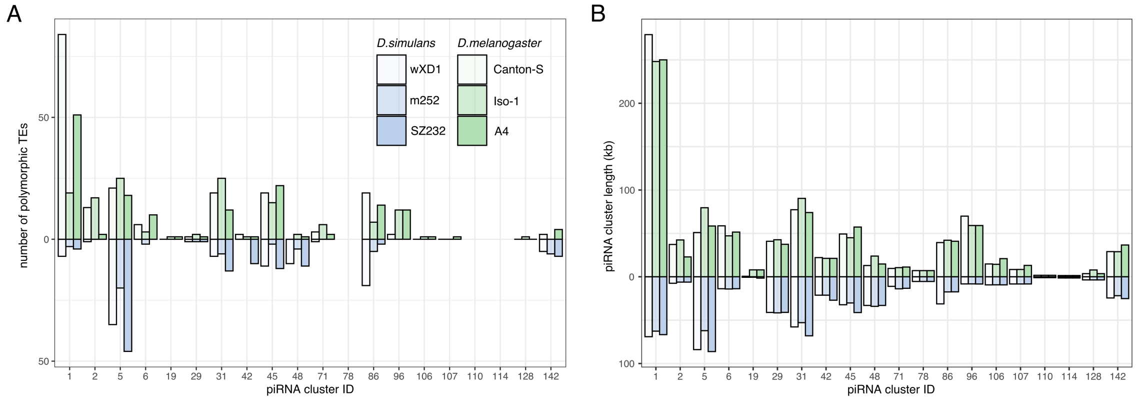


Figure S18: Number of polymorphic TEs in piRNA clusters (A) and length of piRNA clusters (B) in different assemblies of *D. melanogaster* and *D. simulans*

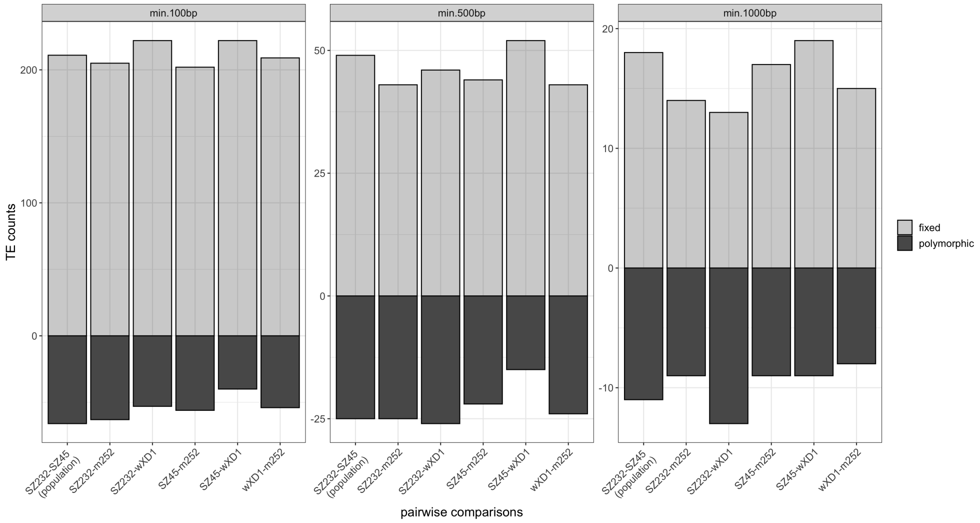


Figure S19: Overview of fixed and polymorphic TE insertions in piRNA clusters based on pairwise comparisons of different *D. simulans* strains. Results are shown for different minimum sizes of TE insertions (top panel).

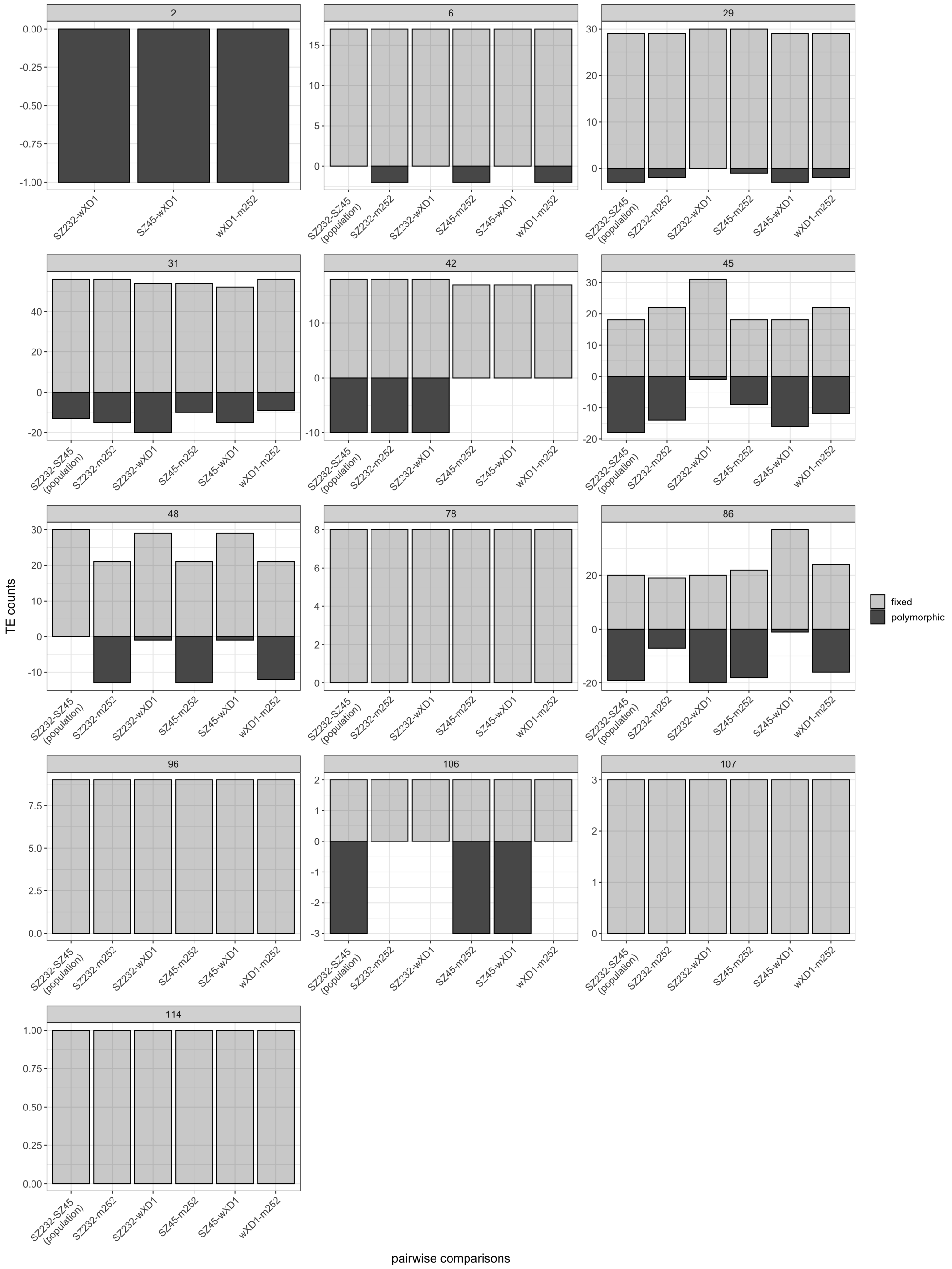


Figure S20: Overview of fixed and polymorphic TE insertions in piRNA clusters based on pairwise comparisons of different *D. simulans* strains. Solely clusters with at least one insertion are shown (minimum size 100bp).

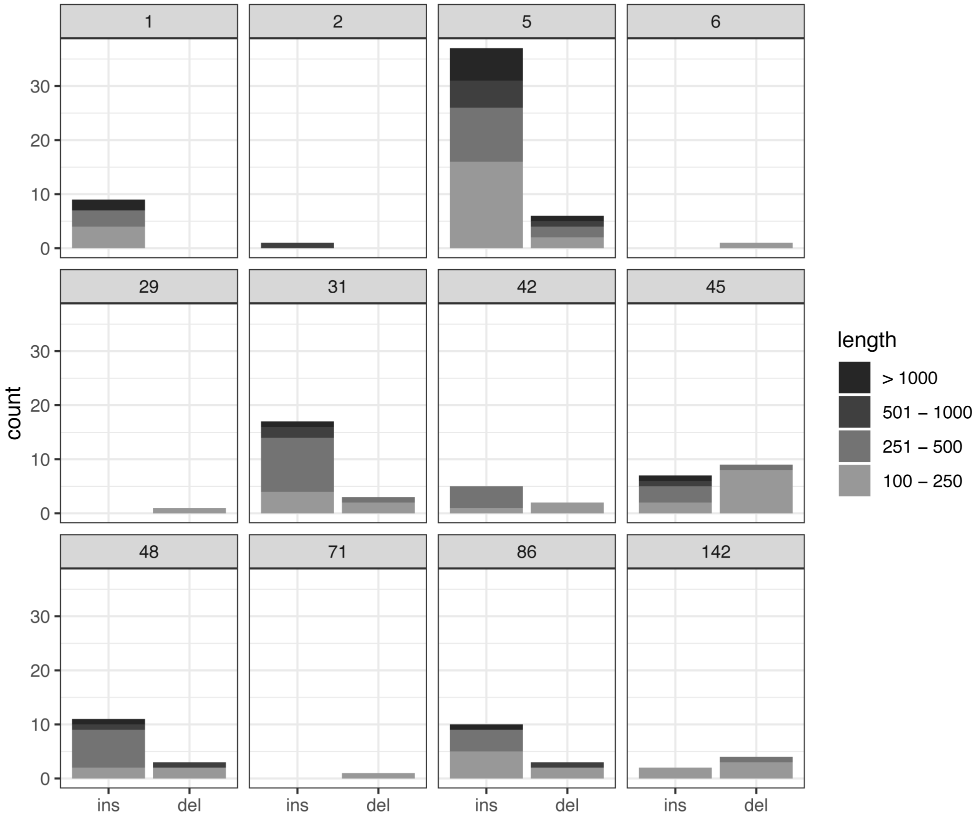


Figure S21: Overview of insertions (ins) and deletions (del) in piRNA clusters of *D. simulans*. Solely clusters with at least one indel are shown. We used the clusters of *D. mauritiana* to polarize the indels. The minimum size of indels was 100bp.

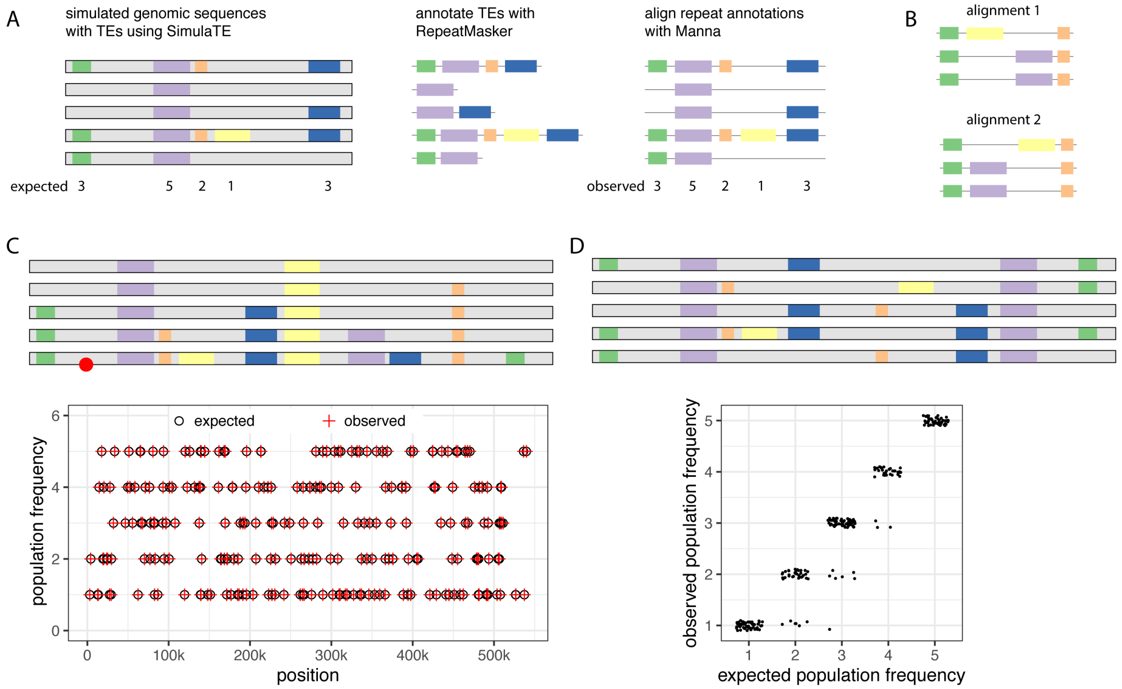


Figure S22: Validating our approach for comparing the composition of piRNA clusters. A) Overview of our validation approach. We simulated a piRNA cluster for five individuals in a population (grey bars) and introduced TE insertions (each color represents a TE family) with a random position and population frequency into the cluster. TEs in the cluster were annotated with RepeatMasker and the annotations were aligned with Manna, enabling us to compare the expected and the observed populating frequency of the insertions. B) For some alignments the positions of gaps, and thus the order of the TEs in the alignment, may be arbitrary. This could cause problems in our validations as in some cases the expected population frequency may not be unambiguously inferred for the observed TE insertions. C) To avoid ambiguous ordering of the TEs we first simulated a scenario where one individual (marked red) carries all 246 simulated TE insertions (two for each of 123 TE families). The bottom figure illustrates that our approach correctly reproduces the expected order and population frequency of the 246 TE insertions. D) Finally we simulated a more challenging scenario with 246 random insertions, where each haplotype solely carries some of the 246 insertions. The ordering of the TEs may be ambiguous in this scenario. In this scenario both insertions of a family had the same population frequency which allowed us to infer the expected population frequency for each TE insertion in the alignment. The jitter plot at the bottom shows that expected and observed population frequencies largely agree.

### Supplementary results; Validating our approach for comparing the composition of piRNA clusters

We thoroughly validated our approach for comparing the composition of piRNA clusters with simulated genomes (supplementary fig. S22). We simulated populations consisting of five haploid genomes (fig. S22A, grey). TE insertions with different population frequencies were introduced into the naive genomes using SimulaTE [Kofler et al., 2018]. Next we annotated TEs in these genomes using RepeatMasker [Smit et al., 1996-2010], aligned the annotations using Manna (supplementary fig. S22A) and tested whether the observed and expected population frequencies of the TE insertions agree. To avoid mismatches between different TE families we set the gap penalty to a lower value than the mismatch penalty. As a consequence the position of gaps and thus the ordering of the TE in the alignment will be arbitrary in some cases, for example when different TE families are inserted into orthologous regions of the aligned strains (supplementary fig. S22B). Ambiguous ordering may be a problem in the validation as it will make it difficult to infer the expected population frequency for the observed TE insertions in the alignment. We first simulated five genomes with 246 TE insertions (2 for each of 123 TE families found in *D. melanogaster*) with a random population frequency between $1-5$ (supplementary fig. S22C). To avoid ambiguous ordering of the TE insertions, one genome contained each of the 246 TE insertions (fig. S22C, marked by red dot). Our approach accurately reproduced the ordering as well as the population frequency of each TE insertion (supplementary fig. S22C). Finally we validated our approach with a challenging scenario, where, in addition to the position and the frequency also the haplotype was randomly selected (supplementary fig. S22D). As a consequence no individual had all 246 TE insertions (two of each family). Both insertions of a given TE family had the same population frequency, which allows to unambiguously infer the expected population frequency for any TE insertion in the alignment (i.e. expectations are based on the identity of the family and not the order of the insertions; supplementary fig. S22D). We correctly estimated the population frequency for the vast majority of the TE insertions (230 out of 246; supplementary fig. S22D). For 16 TE insertions the population frequency was slightly underestimated (supplementary fig. S22D). Several more validations of Manna and the code to reproduce the validations are available at `https://sourceforge.net/p/manna/wiki/Home/#validation`.

#### Supplementary tables

Table S1: Overview of our reciprocal mapping approach to support homology of the 20 piRNA clusters shared between *D. melanogaster* and *D. simulans*. We first aligned the previously published sequences flanking the *D. melanogaster* piRNA clusters to the *D. simulans* genome ($w^{xD1}$), which allows us to identify the positions of these cluster in *D. simulans* (chr., f1.pos, f2.pos). Next, we designed sequences with a length of 1kb flanking these piRNA clusters in *D. simulans* (positions: rf1.ori, rf2.ori) and aligned them to the release 5 of *D. melanogaster* reference genome (Iso-1) allowing us to identify the positions of the flanking sequences in *D. melanogaster* (chr, rf1.pos, rf2.pos). Finally, we ascertained that the coordinates of the piRNA clusters in *D. melanogaster* (chr, c.start, c.end; [Brennecke et al., 2007]) are between the positions of the flanking sequences (i.e. $rf1.pos \leq c.start \leq c.end \leq rf2.pos$). Reverse complemented clusters (compared to [Brennecke et al., 2007]) are shown bold.

| | *D. simulans* | | | | | *D. melanogaster* | | | | |
|---|---|---|---|---|---|---|---|---|---|---|
| c.id | chr. | rf1.ori | f1.pos | f2.pos | rf2.ori | chr. | rf1.pos | c.start | c.end | rf2.pos |
| 1 | 2R | 4,661,253 | 4,662,255 | 4,731,446 | 4,732,096 | 2R | 2,141,639 | 2,144,349 | 2,386,719 | 2,391,296 |
| 2 | X | 21,016,772 | 21,017,774 | 21,026,845 | 21,027,307 | X | 21,387,604 | 21,392,175 | 21,431,907 | 21,433,521 |
| 5 | 2L | 20,044,115 | 20,045,117 | 20,129,357 | 20,129,977 | 2L | 20,146,745 | 20,148,259 | 20,227,581 | 20,228,243 |
| 6 | 3L | 22,866,130 | 22,867,132 | 22,881,160 | 22,881,342 | 3L | 23,271,141 | 23,273,964 | 23,314,199 | 23,319,701 |
| 19 | X | 10,780,896 | 10,781,898 | 10,787,247 | 10,787,586 | X | 11,083,358 | 11,089,529 | 11,095,706 | 11,096,773 |
| 29 | 4 | 739,454 | 740,456 | 782,498 | 783,631 | 4 | 822,024 | 824,220 | 865,373 | 867,965 |
| 31 | 2L | 22,864,254 | 22,865,256 | 22,923,869 | 22,925,036 | 2L | 22,476,617 | 22,489,509 | 22,556,902 | 22,569,036 |
| 42 | 3R | 1,178,793 | 1,179,795 | 1,201,815 | 1,201,963 | 3R | 125 | 6,252 | 22,624 | 23,385 |
| **45** | **X** | **22,399,288** | **22,366,575** | **22,398,981** | **22,365,573** | **X** | **22,339,663** | **22,342,541** | **22,383,561** | **22,384,388** |
| 48 | 3L | 19,575,898 | 19,576,900 | 19,611,235 | 19,611,376 | 3L | 19,835,637 | 19,841,657 | 19,861,753 | 19,862,197 |
| 71 | 2R | 3,778,303 | 3,780,305 | 3,791,621 | 3,792,346 | 2R | 1,214,908 | 1,220,249 | 1,225,114 | 1,228,018 |
| 78 | 4 | 922,676 | 923,678 | 929,171 | 929,374 | 4 | 1,017,946 | 1,020,285 | 1,025,978 | 1,026,636 |
| **86** | **3L** | **23,339,071** | **23,307,533** | **23,338,897** | **23,307,651** | **3L** | **24,159,836** | **24,171,077** | **24,199,822** | **24,202,885** |
| **96** | **3R** | **402,380** | **393,214** | **401,971** | **392,212** | **3RHet** | **2,158,207** | **2,178,310** | **2,199,481** | **2,205,043** |
| 106 | 4 | 534,401 | 535,403 | 545,135 | 546,076 | 4 | 608,470 | 612,204 | 623,399 | 625,615 |
| 107 | 2L | 22,000,290 | 22,001,292 | 22,009,900 | 22,010,467 | 2L | 21,894,195 | 21,894,816 | 21,900,308 | 21,904,121 |
| 110 | X | 12,320,899 | 12,321,901 | 12,323,639 | 12,324,171 | X | 12,663,115 | 12,664,996 | 12,666,347 | 12,666,941 |
| 114 | X | 3,824,141 | 3,825,143 | 3,827,177 | 3,827,359 | X | 4,019,648 | 4,022,252 | 4,022,455 | 4,022,880 |
| 128 | 2R | 5,629,535 | 5,630,537 | 5,634,276 | 5,634,466 | 2R | 3,319,269 | 3,321,225 | 3,327,008 | 3,328,474 |
| 142 | 2R | 3,634,403 | 3,636,405 | 3,661,781 | 3,662,354 | 2R | 1,033,509 | 1,039,689 | 1,048,724 | 1,066,362 |

Table S2: Overview of our reciprocal mapping approach to support homology of the 20 piRNA clusters shared between *D. melanogaster* and *D. sechellia*. We first aligned the previously published sequences flanking the *D. melanogaster* piRNA clusters to the *D. sechellia* genome (sech25), which allows us to identify the positions of these cluster in *D. sechellia* (chr., f1.pos, f2.pos). Next, we designed sequences with a length of 1kb flanking these piRNA clusters in *D. sechellia* (positions: rf1.ori, rf2.ori) and aligned them to the release 5 of *D. melanogaster* reference genome (Iso-1) allowing us to identify the positions of the flanking sequences in *D. melanogaster* (chr, rf1.pos, rf2.pos). Finally, we ascertained that the coordinates of the piRNA clusters in *D. melanogaster* (chr, c.start, c.end; [Brennecke et al., 2007]) are between the positions of the flanking sequences (i.e. $rf1.pos \leq c.start \leq c.end \leq rf2.pos$). Reverse complemented clusters (compared to [Brennecke et al., 2007]) are shown bold.

| | *D. sechellia* | | | | | *D. melanogaster* | | | | |
|---|---|---|---|---|---|---|---|---|---|---|
| c.id | chr. | rf1.ori | f1.pos | f2.pos | rf2.ori | chr. | rf1.pos | c.start | c.end | rf2.pos |
| 1 | 2R | 2,712,071 | 2,713,073 | 2,797,604 | 2,798,254 | 2R | 2,141,644 | 2,144,349 | 2,386,719 | 2,391,460 |
| 2 | X | 21,608,301 | 21,609,303 | 21,616,833 | 21,617,295 | X | 21,387,579 | 21,392,175 | 21,431,907 | 21,433,521 |
| 5 | 2L | 19,737,779 | 19,738,781 | 19,790,046 | 19,790,666 | 2L | 20,146,745 | 20,148,259 | 20,227,581 | 20,228,245 |
| 6 | 3L | 22,599,800 | 22,599,553 | 22,618,886 | 22,619,068 | 3L | 23,271,563 | 23,273,964 | 23,314,199 | 23,319,701 |
| 19 | X | 10,967,784 | 10,968,786 | 10,972,467 | 10,972,806 | X | 11,084,324 | 11,089,529 | 11,095,706 | 11,096,773 |
| 29 | 4 | 786,089 | 787,091 | 828,722 | 829,855 | 4 | 822,045 | 824,220 | 865,373 | 867,909 |
| 31 | 2L | 23,390,685 | 23,391,687 | 23,454,063 | 23,455,230 | 2L | 22,476,617 | 22,489,509 | 22,556,902 | 22,569,036 |
| 42 | 3R | 3,073,137 | 3,074,139 | 3,096,518 | 3,096,666 | 3R | 125 | 6,252 | 22,624 | 23,385 |
| **45** | **X** | **22,140,704** | **22,104,512** | **22,140,397** | **22,103,510** | **X** | **22,337,995** | **22,342,541** | **22,383,561** | **22,384,388** |
| 48 | 3L | 19,295,649 | 19,296,651 | 19,313,914 | 19,314,055 | 3L | 19,835,635 | 19,841,657 | 19,861,753 | 19,862,197 |
| 71 | 2R | 1,740,568 | 1,742,570 | 1,759,683 | 1,760,408 | 2R | 1,214,888 | 1,220,249 | 1,225,114 | 1,228,018 |
| 78 | 4 | 981,145 | 982,147 | 987,557 | 987,760 | 4 | 1,017,946 | 1,020,285 | 1,025,978 | 1,026,636 |
| **86** | **3L** | **23,725,708** | **23,704,712** | **23,725,534** | **23,704,830** | **3L** | **24,159,836** | **24,171,077** | **24,199,822** | **24,202,910** |
| 96 | 3R | 2,210,457 | 2,211,459 | 2,276,313 | 2,276,845 | 3RHet | 2,158,207 | 2,178,310 | 2,199,481 | 2,205,161 |
| 106 | 4 | 588,340 | 589,342 | 603,960 | 604,901 | 4 | 608,330 | 612,204 | 623,399 | 625,614 |
| 107 | 2L | 22,169,195 | 22,170,197 | 22,178,586 | 22,179,153 | 2L | 21,894,195 | 21,894,816 | 21,900,308 | 21,904,121 |
| 110 | X | 12,535,122 | 12,536,124 | 12,537,865 | 12,538,397 | X | 12,663,115 | 12,664,996 | 12,666,347 | 12,667,067 |
| 114 | X | 3,903,460 | 3,904,462 | 3,906,548 | 3,906,730 | X | 4,019,641 | 4,022,252 | 4,022,455 | 4,022,880 |
| 128 | 2R | 3,703,212 | 3,704,214 | 3,708,063 | 3,708,253 | 2R | 3,319,268 | 3,321,225 | 3,327,008 | 3,328,474 |
| 142 | 2R | 1,568,770 | 1,571,772 | 1,608,533 | 1,611,106 | 2R | 1,032,090 | 1,039,689 | 1,048,724 | 1,067,370 |

Table S3: Overview of our reciprocal mapping approach to support homology of the 20 piRNA clusters shared between *D. melanogaster* and *D. mauritiana*. We first aligned the previously published sequences flanking the *D. melanogaster* piRNA clusters to the *D. mauritiana* genome (mau12), which allows us to identify the positions of these cluster in *D. mauritiana* (chr., f1.pos, f2.pos). Next, we designed sequences with a length of 1kb flanking these piRNA clusters in *D. mauritiana* (positions: rf1.ori, rf2.ori) and aligned them to the release 5 of *D. melanogaster* reference genome (Iso-1) allowing us to identify the positions of the flanking sequences in *D. melanogaster* (chr, rf1.pos, rf2.pos). Finally, we ascertained that the coordinates of the piRNA clusters in *D. melanogaster* (chr, c.start, c.end; [Brennecke et al., 2007]) are between the positions of the flanking sequences (i.e. $rf1.pos \leq c.start \leq c.end \leq rf2.pos$). Reverse complemented clusters (compared to [Brennecke et al., 2007]) are shown bold.

| | *D. mauritiana* | | | | | *D. melanogaster* | | | | |
|---|---|---|---|---|---|---|---|---|---|---|
| c.id | chr. | rf1.ori | f1.pos | f2.pos | rf2.ori | chr. | rf1.pos | c.start | c.end | rf2.pos |
| 1 | 2R | 5,004,999 | 5,006,001 | 5,054,026 | 5,054,676 | 2R | 2,141,633 | 2,144,349 | 2,386,719 | 2,391,296 |
| 2 | X | 21,296,756 | 21,297,758 | 21,305,432 | 21,305,894 | X | 21,387,606 | 2,139,2175 | 21,431,907 | 21,433,521 |
| 5 | 2L | 19,669,352 | 19,670,354 | 19,709,473 | 19,710,093 | 2L | 20,146,749 | 20,148,259 | 20,227,581 | 20,228,241 |
| 6 | 3L | 22,572,463 | 22,573,465 | 22,589,904 | 22,590,086 | 3L | 23,271,135 | 23,273,964 | 23,314,199 | 23,319,708 |
| 19 | X | 10,966,963 | 10,968,965 | 10,972,078 | 10,972,417 | X | 11,083,889 | 11,089,529 | 11,095,706 | 11,096,773 |
| 29 | 4 | 716,446 | 717,448 | 759,689 | 760,822 | 4 | 822,024 | 824,220 | 865,373 | 868,052 |
| 31 | 2L | 22,609,202 | 22,610,204 | 22,668,741 | 22,667,739 | 2L | 22,476,617 | 22,489,509 | 22,556,902 | 22,568,531 |
| 42 | 3R | 1,567,943 | 1,568,945 | 1,591,138 | 1,591,286 | 3R | 125 | 6,252 | 22,624 | 23,385 |
| **45** | **X** | **22,438,911** | **22,397,556** | **22,438,604** | **22,396,554** | **X** | **22,339,663** | **22,342,541** | **22,383,561** | **22,384,388** |
| 48 | 3L | 19,299,452 | 19,300,454 | 19,318,721 | 19,318,862 | 3L | 19,835,636 | 19,841,657 | 19,861,753 | 19,862,197 |
| 71 | 2R | 4,040,929 | 4,041,931 | 4,055,698 | 4,056,423 | 2R | 1,215,978 | 1,220,249 | 1,225,114 | 1,228,018 |
| 78 | 4 | 910,025 | 911,027 | 916,846 | 917,049 | 4 | 1,018,308 | 1,020,285 | 1,025,978 | 1,026,636 |
| **86** | **3L** | **23,058,759** | **23,035,112** | **23,058,585** | **23,037,230** | **3L** | **24,159,836** | **24,171,077** | **24,199,822** | **24,202,153** |
| **96** | **3R** | **1,058,992** | **1,047,072** | **1,058,583** | **1,046,070** | **3RHet** | **2,158,207** | **2,178,310** | **2,199,481** | **2,205,043** |
| 106 | 4 | 517,394 | 518,396 | 527,294 | 528,235 | 4 | 608,330 | 612,204 | 623,399 | 625,614 |
| 107 | 2L | 21,478,245 | 21,479,247 | 21,492,793 | 21,493,360 | 2L | 21,894,195 | 21,894,816 | 21,900,308 | 21,904,121 |
| 110 | X | 12,567,267 | 12,568,269 | 12,570,008 | 12,570,540 | X | 12,663,115 | 12,664,996 | 12,666,347 | 12,666,941 |
| 114 | X | 3,965,780 | 3,966,782 | 3,968,881 | 3,969,063 | X | 4,019,641 | 4,022,252 | 4,022,455 | 4,022,880 |
| 128 | 2R | 5,966,549 | 5,967,551 | 5,971,338 | 5,971,528 | 2R | 3,319,305 | 3,321,225 | 3,327,008 | 3,328,474 |
| 142 | 2R | 3,876,829 | 3,878,831 | 3,913,130 | 3,913,703 | 2R | 1,035,210 | 1,039,689 | 1,048,724 | 1,066,362 |

Table S4: Similarity of the piRNA clusters within and between species. The similarity reflects the average fraction of TE sequences that can be aligned between two orthologous clusters (%). Values are the average similarity among all possible pairwise comparisons (e.g. three for *D. melanogaster*: A4 vs. Iso-1, A4 vs. Canton-S, Iso-1 vs. Canton-S). cl.: cluster, w.av.: weighted average for all clusters

| cl. | mel | sim | mel-sim | sec-sim | mau-sim |
|---|---|---|---|---|---|
| 1 | 77.5 | 90.2 | 4 | 26.4 | 29.6 |
| 2 | 45.4 | 33.3 | 0.4 | 0 | 66.7 |
| 5 | 59.1 | 51.5 | 10.4 | 16.7 | 15.9 |
| 6 | 83.8 | 96.8 | 9.8 | 48.8 | 70.6 |
| 19 | 33.3 | 0 | 0 | 0 | 0 |
| 29 | 89.5 | 96.3 | 36.7 | 80.3 | 81.8 |
| 31 | 73 | 75.8 | 3.4 | 42.5 | 53.5 |
| 42 | 93.6 | 72.7 | 41.5 | 79.4 | 73 |
| 45 | 60.1 | 60.1 | 7.7 | 42.5 | 42.8 |
| 48 | 54.3 | 74.7 | 26.7 | 21.3 | 29 |
| 71 | 88.1 | 93.6 | 2.2 | 53.1 | 68.8 |
| 78 | 100 | 99.8 | 47.2 | 73.5 | 84.6 |
| 86 | 66.5 | 63.6 | 3.1 | 47 | 51.4 |
| 96 | 86.4 | 100 | 2 | 6.2 | 59.3 |
| 106 | 62.4 | 100 | 3 | 44.4 | 99.8 |
| 107 | 43.5 | 100 | 60.1 | 69 | 17.7 |
| 110 | 100 | 0 | 0 | 0 | 0 |
| 114 | 100 | 100 | 93.5 | 88.8 | 88.9 |
| 128 | 33.3 | 100 | 47.1 | 59.5 | 100 |
| 142 | 83.6 | 87.5 | 11.9 | 55.4 | 56 |
| w.av. | 73.1 | 74.7 | 8.1 | 32.7 | 41.4 |

Table S5: Similarity of the piRNA clusters within *D. simulans*. The similarity reflects the average fraction of TE sequences that can be aligned between two orthologous clusters (%). cl.: cluster w.av.: weighted average for all clusters

| cl. | SZ232-SZ45 | SZ232-m252 | SZ45-m252 | SZ232-$w^{xD1}$ | SZ45-$w^{xD1}$ | $w^{xD1}$-m252 |
|---|---|---|---|---|---|---|
| 2 | 100 | 100 | 100 | 0 | 0 | 0 |
| 6 | 99.8 | 95.2 | 95.5 | 99.8 | 100 | 95.5 |
| 12 | 81.3 | 87.8 | 74.8 | 80.9 | 73.6 | 83.5 |
| 19 | 0 | 0 | 0 | 0 | 0 | 0 |
| 29 | 81.3 | 94.1 | 84.7 | 98 | 82 | 96.7 |
| 31 | 73.3 | 73.6 | 78 | 70 | 73.3 | 83.6 |
| 42 | 59.5 | 58.6 | 99.4 | 59.4 | 99.4 | 100 |
| 45 | 35.6 | 44.5 | 57.1 | 71.5 | 47.6 | 64.3 |
| 48 | 100 | 62.5 | 62.6 | 98.9 | 98.8 | 62.6 |
| 78 | 99.8 | 99.7 | 100 | 99.8 | 100 | 100 |
| 86 | 51.5 | 87.9 | 50.9 | 51 | 99 | 51.8 |
| 96 | 100 | 100 | 100 | 100 | 100 | 100 |
| 106 | 21.9 | 100 | 21.9 | 100 | 21.9 | 100 |
| 107 | 100 | 100 | 100 | 100 | 100 | 100 |
| 110 | 0 | 0 | 0 | 0 | 0 | 0 |
| 114 | 100 | 100 | 100 | 100 | 100 | 100 |
| 128 | 59.5 | 100 | 59.5 | 100 | 59.5 | 100 |
| w.av. | 72.5 | 73.4 | 71.3 | 76.6 | 81.9 | 75.7 |
